# Ecosystem Composition Controls the Early Fate of Rare Earth Elements during Incipient Soil Genesis

**DOI:** 10.1101/061846

**Authors:** Dragos G. Zaharescu, Carmen I. Burghelea, Katerina Dontsova, Jennifer K. Presler, Raina M. Maier, Travis Huxman, Kenneth J. Domanik, Edward A. Hunt, Mary K. Amistadi, Emily E. Gaddis, Maria A. Palacios-Menendez, Maria O. Vaquera-Ibarra, Jonathan Chorover

## Abstract

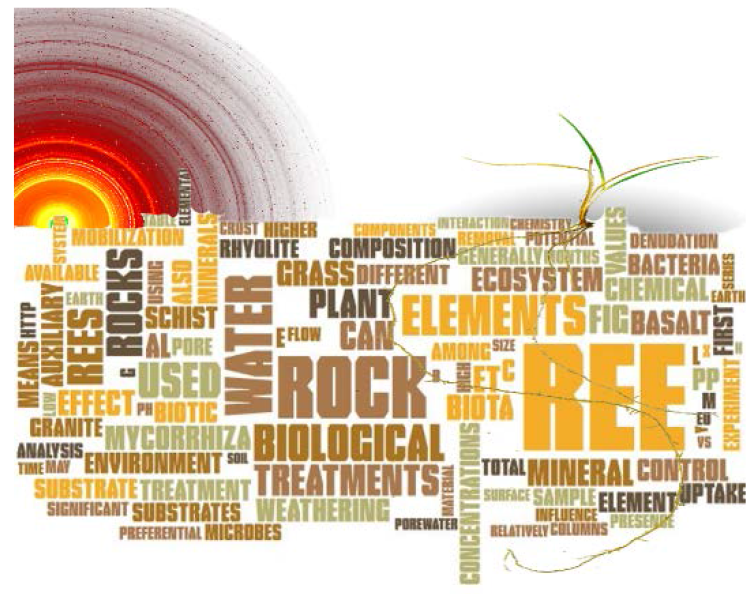

**We used a model rock-biota system to demonstrate that ecosystem composition (microbial and plant) regulates the fate of REE during early biota-rock interactions**.

*Graphical abstract:* word cloud with article keywords, Buffalograss, and X-ray diffractogram.

The rare earth elements (REE) are of increasing importance in a variety of science and economic fields, including (bio)geosciences, paleoecology, astrobiology, and mining. Despite their great promise, REE fractionation in early plant-microbe-rock systems has largely remained elusive. We tested the hypothesis that REE mass-partitioning during the incipient weathering of basalt, rhyolite, granite and schist depends on the activity of microbes, plant, and arbuscular mycorrhiza.

Pore-water element abundances reflected a rapid transition from abiotic to biotic weathering, the latter associated with lower aqueous loss and higher uptake. Abiotic dissolution contributed 38.6±19% to total denudation. Microbes incremented denudation, particularly in rhyolite, this effect associating with decreased bioavailable solid fractions in this rock. Total mobilization (aqueous+uptake) was ten times greater in planted treatments compared to abiotic control, REE masses in plant generally exceeding those in water. Plants of larger biomass further increased solid fractions, consistent with soil genesis. Mycorrhiza had a generally positive effect on total mobilization. The incipient REE weathering was dominated by inorganic dissolution enhanced by biotic respiration, the patterns of denudation largely dictated by mineralogy. A consistent biotic signature was observed in La:phosphate, mobilization:solid fraction in all rocks, as well as in the general pattern of denudation and uptake.

---

The transformation of rock to soil supports Earth’s terrestrial life. Soil genesis and evolution have been under scientific scrutiny for over a century. With recent advances in analytical chemistry, molecular/evolutionary biology, hydrology, ecology and remote sensing we are only now beginning to understand how different components of geosphere, hydrosphere, atmosphere, and biosphere work together at different scales to shape the surface of Earth and transform parent rock into soil that sustains the ecosystem^1,22,3,1,4,5^. Among the most reactive phases in the transformation of crustal rock are the earliest stages of mineral weathering when first microbial and plant communities interact with bedrock and spark the flow of energy and nutrients that feed into major biogeochemical cycles. The igneous rock-Earth’s nutrient store, exhibits its largest thermodynamic disequilibrium at surface pressure and temperatures, where the oxic aqueous environment makes it highly susceptible to weathering. Rapid mineral transformations of crustal rock often accompany the earliest stages of weathering, and they are intensified under mineral colonization by microbiota and plant roots. Studying these early interactions in the natural soil-forming system has classically been focused on nutrients that are part of plant and microorganism metabolic budgets. These studies have shown, for example, preferential dissolution/ loss of major elements such as Fe, Na and Ca from micaceous minerals and plagioclase^6,7^, microbe and plant elemental uptake, and incorporation into newly formed minerals^8,9^. Resulting biological signals of these elements in the complex porous geomedia can, however, be masked by their high and diverse reactivity, and competing chemical processes. The use of less mobile elements as indicators of biological weathering is potentially more powerful because of their simpler cycle and comparatively stronger selective force organisms would need to affect their stoichiometry.

Rare earth elements (REE) are the lanthanide series (atomic numbers [Z] 57 [lanthanum] to 71 [lutetium]) with yttrium (Z=39) often included because its outer electron shell structure and ionic radius are nearly identical to holmium^10^. REE exhibit generally similar but highly dispersed environmental distributions. Coherent trends in their aqueous reactivity derive from similar stable arrangement of outer electron configuration (5*sp*) across the series, superimposed with ionic radius variation resulting from gradual filling of the inner 4f electron shell. The lower polarization of empty (La, Y), half-filled (Gd) and filled (Lu) 4f electron shells, as well as variable redox states of Ce and Eu can lead to radius-independent fractionation behavior^11^. The series exhibits a consistent decrease in ionic radius with increasing atomic mass (lanthanide contraction effect). This decrease in ionic radius with increasing atomic number along with slight differences in ionic potential and unpaired 4f electrons can induce variability in REE chemical behavior^12,13^. Some fractionation in low temperature natural terrestrial systems is therefore possible, particularly at circumneutral pH^14^, due to differential weathering of minerals^15^; surface adsorption reactions of dissolved REE^16^, co-precipitation in secondary minerals (e.g. silicate clays and Mn-, Al-, Fe-(oxy)hydroxides^17^, competition with major ions on organic binding sites^18^ and soluble complex formation with a variety of ligands, including carbonates and humic acids^15^. Likewise, sub-partition smoothed curves along REE series (tetrad effect) have been described in shale-normalized REE series, indicating their involvement in ionic radius/charge independent processes^19^. The tetrad effect is presumably related to the increased stability at quarter, half, three-quarter, and complete filling of the 4f electron shell^20^. Systematic variation in REE reactivity across the series, uncommon to other elements, has stimulated their use as tracers of a variety of geochemical processes, from mantle and crustal to cosmogenic evolution, ore genesis, sedimentary petrology^21^, water-rock interactions^22^, and critical zone evolution^23^.

We postulate that since REE form strong complexes with bio-ligands^23,24^, biological ecosystem components (microbes and plants) can affect the behavior of REE during weathering, e.g., by selective ligand-complexation of dissolved REE ions. REE cycles in the biosphere are still poorly understood, with conflicting evidence and opinions regarding biological effects on geochemical cycles and the role of REE in biological systems. In regard to the latter, suggested interactions include: REE stimulation of plant biomass production^12^, disruption of physiological functions, e.g. floral development and photosynthesis by replacing major functional metals such as Ca in a nutrient-deprived environment^25,26^, and biosorption by microorganisms^27^. Moreover, arbuscular mycorrhiza, the most widespread type of soil fungal-plant symbiosis associated with 74% of flowering plant species^28^, can both, reduce and increase the extraction of REE ions from mine tailings^29^. Adding to this debate is a report of lanthanides being essential for some acidophilic methanotrophic microbes by providing superior catalytic properties to dehydrogenase proteins^30^. However, biological effects on incipient REE cycles in natural conditions are practically unknown.

While fractionation patterns of REE in mineral weathering environments has the potential to provide signals of processes leading to mineral dissolution and secondary mineral formation^31^, much less is known about the role of this class of elements in the incipient biological transformation of Earth’s crust. Hypothetically, due to their similar mineral geochemistry and their reactivity with diverse biological ligands, REE could be suitable long term indicators for a wider range of biological alteration of rock surfaces better than other elements. Knowledge of their behavior in such a context may provide signatures of life’s presence and the extent of its influence on bedrock on early and modern Earth, and potentially other planetary bodies.

Here we present results from a highly controlled experiment designed to quantify the extent to which variation in the nature of incipient bio-weathering results in the mobilization and redistribution of REE during silicate (igneous and metamorphic) rock weathering. We hypothesized that rock type and the nature of rock-colonizing biotic communities would affect REE fractionation and that differential and reproducible fractionation patterns would, therefore, be observed in pore waters, biological tissue, and solid-phase extractable pools. Specifically, we tested whether: (a) rock substrate controls both chemical and biological REE mobilization in pore water; (b) there is a distinct signal from biotic presence on pore water REE content; (c) there is selective uptake and distribution of REE in plants, which (d) is affected by the presence of arbuscular mycorrhizal fungi; and (e) there is a biotic influence on the soluble and poorly crystalline mineral pools.

**Figure 1.**
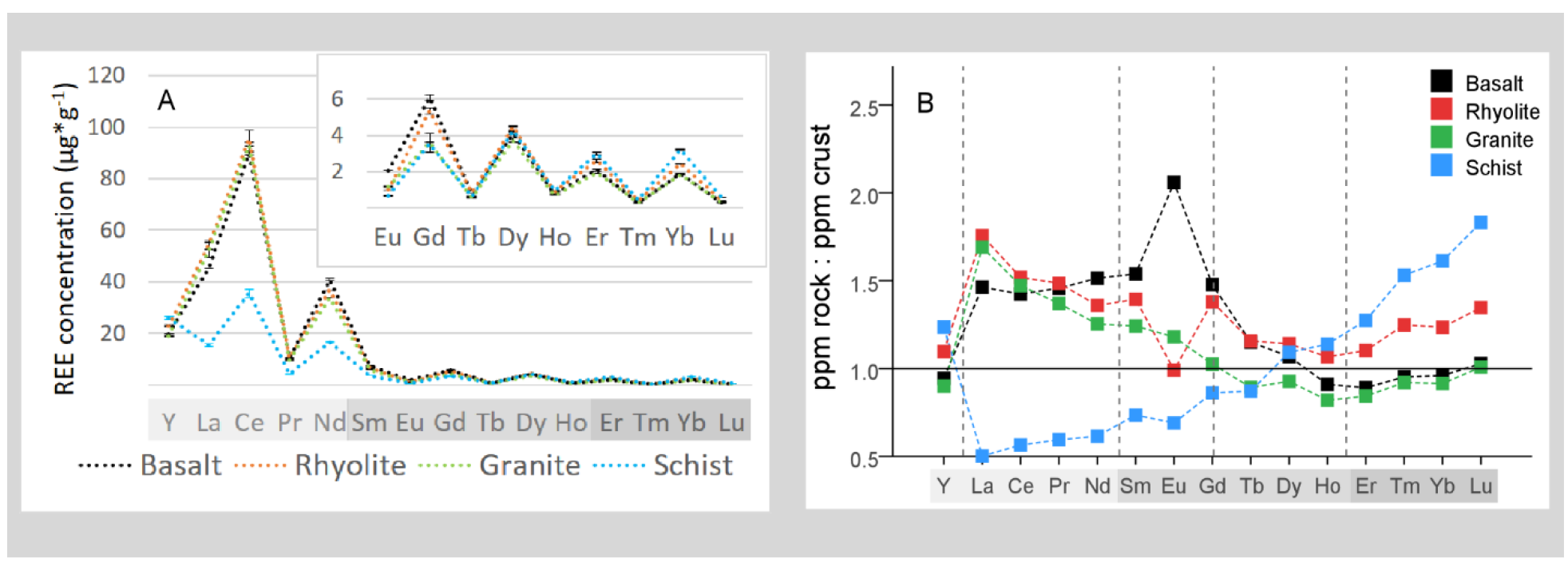
REE abundances in the original rock. (A) REE concentration in bulk rocks. (B) The concentration of REE in the initial rock substrates normalized to Earth’s upper continental crust averages ^32^. Tetrad groups (except Y) are separated by vertical dashed lines. Elements are arranged in increasing atomic number with those defined as low (L), medium (M) and heavy (H) atomic mass displayed on X axis with light, medium and dark gray background, respectively.

## Results and discussion

**Rock substrates and REE sources.** Initial concentrations of various REE in the four rocks were similar except for schist, which had less light (L)-REE and more heavy (H)-REE than the other rocks (Figure 1A). Schist was also depleted in L-REE relative to upper continental crust while basalt, rhyolite, and granite were enriched (Figure 1B). A weak M-class tetrad effect ^19^ was observed in granite (Figure 1B). Europium exhibited a positive anomaly in basalt and a negative one in rhyolite. In basalt, microprobe analysis identified Ce-Nd-La-Pr oxides as mineral hosts for L-REE, and the glass matrix as the source of heavier REE (Fig. s3). Principal hosts in rhyolite included ilmenite (rich in La, Gd, and Yb) and apatite (rich in Y). In granite, sphene, apatite (both rich in L-REE) and K-feldspars were predominant hosts, whereas in schist, zircon (rich in Yb), allanite (rich in La) and xenotime (rich in Y) were prevalent. These differences in bedrock chemistry and mineralogy are expected to influence the stoichiometry of REE release during weathering.

**Denudation of REE in solution.** When exposed to water, all rocks exhibited rapid REE release to aqueous solution (i.e., chemical denudation), which slowed after initial two months (Figure 2). Microbes and plants induced greater denudation than the control in basalt and rhyolite, however, the presence of arbuscular mycorrhiza increased denudation with respect to plant-microbe in basalt and schist and reduced it in rhyolite probably related differential retention in pore space. For granite, bacteria alone had no effect; only planted treatments were significantly different from control. Also in schist, bacteria and buffalo grass treatments were lower than control. A significant divergence between treatments, where present, started to develop in pore water within 4-6 weeks of seeding and/or inoculation (Figure 2).

**Figure 2.**
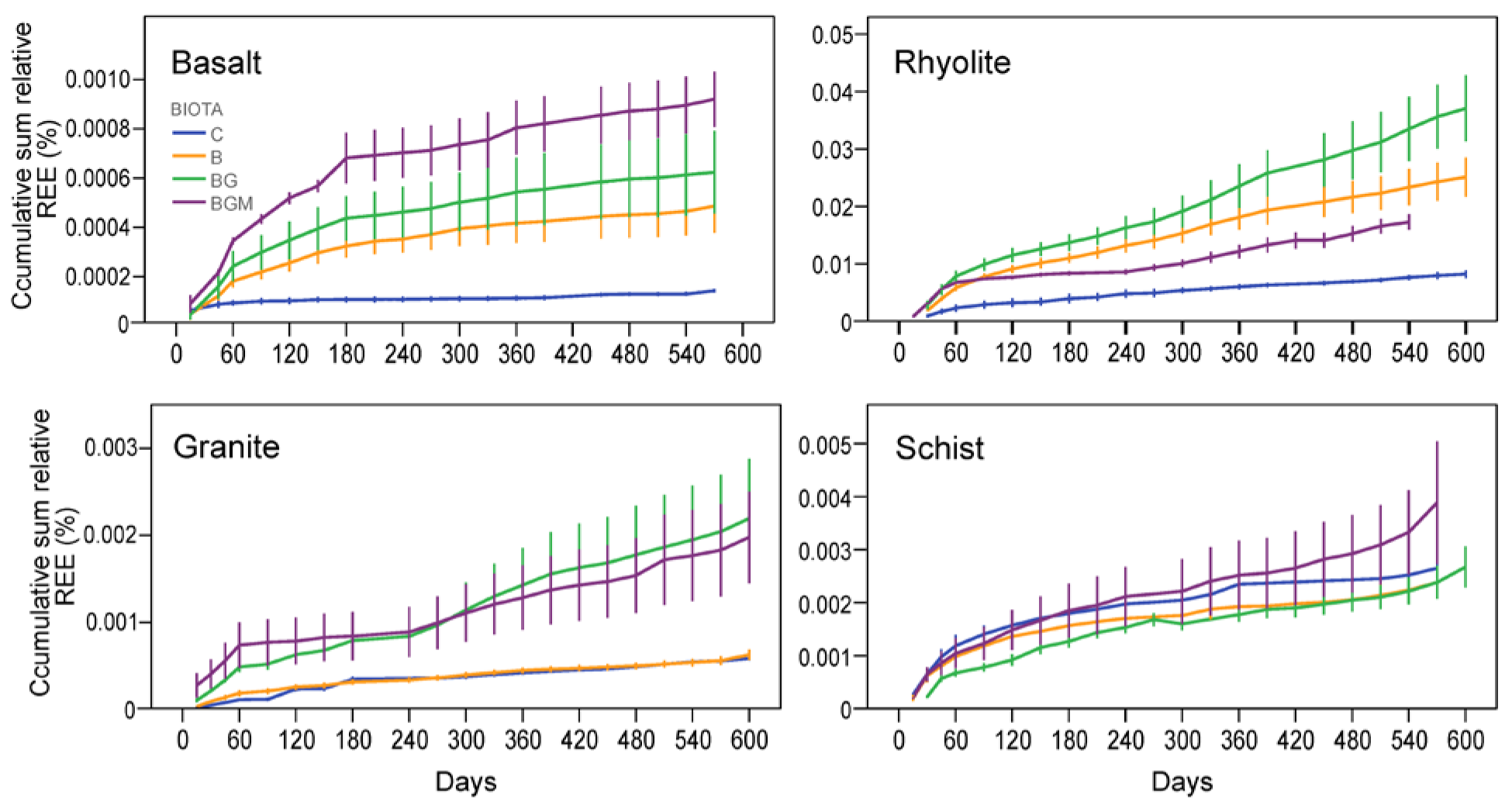
REE preferential denudation in time under abiotic and biotic influence. Cumulative denudation per column for a sum of all REE relative to their rock content. Values are means of three replicate columns and error bars represent one standard error of the mean. C, control; B, bacteria; BG, bacteria-grass; BGM, bacteria-grass-mycorrhiza.

The observed pattern of denudation is consistent with a two-phase process: an initial highly reactive phase, when readily available sites on fresh mineral surfaces release REE in pore space through mostly abiotic water-rock interactions, e.g. hydrolysis or carbonation; followed by a phase with slower increase in denudation but greater biological influence. Previous observations of major element (Si, Ca and K) leaching from materials of comparable particle size and mineralogy have shown a high initial removal in biota and biota-free substrates followed by a steady-state phase^33^. This is consistent with our measured changes in root growth; in the first 120 days, roots accumulated 0.12±0.06g of biomass per column, or 1198±327cm total root length, while at the end of the experiment (~600 days) they measured 0.14±0.04g and 2728±2577cm, indicating that majority of plant growth happened during the first months of biotic establishment.

**Figure 3.**
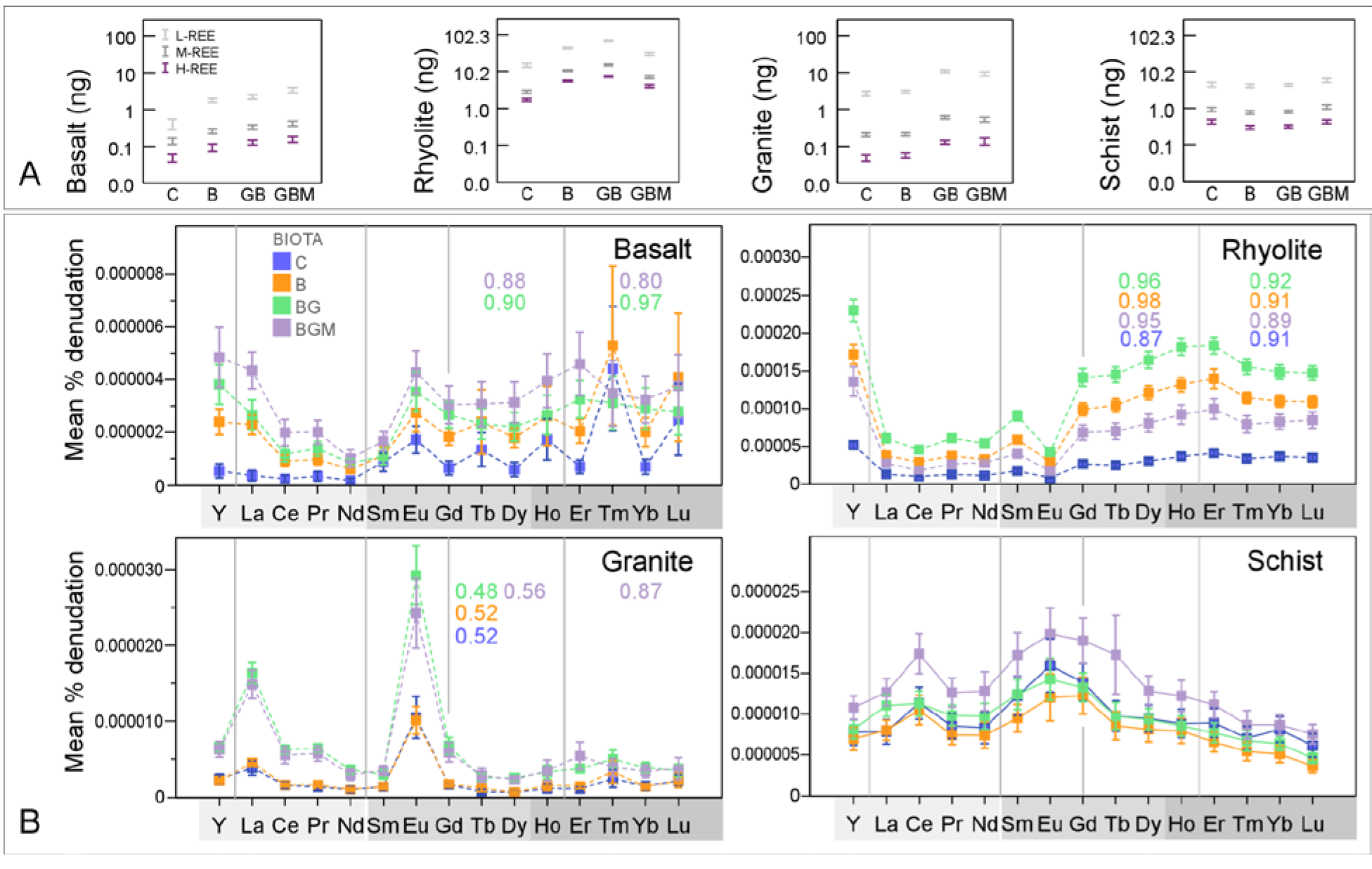
Abiotic and biotic influence REE variability in pore-water. (A) Total denudation of L-, M-and H-REE in pore water collected over 20 months experiment for following treatments: C, control; B, bacteria; BG, bacteria-grass; BGM, bacteria-grass-mycorrhiza. (B) Rock normalized (g in column: g in column, %) sample means of L-, M-and H-REE in pore-water as a function of rock and ecosystem composition. Error bars represent one standard error (SE) of the mean. Tetrad groups (Y excluded) are separated by vertical lines. Numbers inset represent tetrad magnitude values (numbers range from 1-no tetrad to higher and lower values for M-type and W-type curves), with color corresponding to treatment color.

Total REE mass removed in solution differed by rock (Table s2). Rhyolite showed the highest REE denudation followed by schist, granite, and basalt. Biological treatment effects observed for REE were also observed for other water parameters, including electrical conductivity (EC). For example, Basalt, which showed the greatest REE denudation for planted treatments relative to control also had higher EC in planted treatments, indicating greater weathering overall. Other potential indicators of weathering, such as solution pH and total organic carbon were less sensitive to the biological effect.

Trends observed for the sum of REE (Figure 2) were also observed for groups of REE (Figure 3A) and for individual elements (Figure 3B). Among elements, L-REE were subjected to greatest denudation followed by M-REE and H-REE (Figure 3A), consistent with their rock abundances (Figure 1A). Rock-normalized REE concentrations revealed similar trends with the time-lapse analysis, with the notable exception of schist where the biotic effect of vascular plants (BG) over microbes (B) and mycorrhiza (BGM) over BG was clearer, particularly for L-REE (Figure 3B). These differences between biological treatments with substrate imply an important rock-dependent biological effect on REE released from rock during weathering.

A major depression occurred in the L-REE segment in basalt and rhyolite, which was enhanced by biological treatments (Figure 3B). Since both rocks were relatively rich in these elements (Figure 1), the depressions could be caused by comparatively slower dissolution kinetics of hosting minerals and higher uptake. For instance, in basalt, previous studies showed that amorphous glass – the identified source of H-REE in our experiment (Section 2.1 and Fig. s3) is among the first constituents to dissolve^33,34^. This would release higher amounts of H-REE in solution, producing a depression in L-REE. In granite, preferential dissolution of apatite^35^ – targeted by plant and mycorrhiza due to its rich P content^36^ is likely responsible for relatively high L-REE values observed in water collected from this rock (Figure 3B).

Europium and cerium are known to exhibit anomalies due to their variable oxidation states (Eu fractionating during precipitation from magma, and Ce during low-temperature water-mineral interaction). Europium anomalies were recorded in basalt, granite (even though no Eu anomaly was present in parent granite; Figure 1B) and rhyolite, and Ce anomaly in schist (Figure 3B). Europium fractionation is not uncommon during aqueous weathering of felsic rocks, particularly under organic acids^37^, and can be attributed to Eu preferential release from feldspars and apatite, where Eu is abundant Ca^2+^ substitute (as Eu^2+^)^12^. Cerium – the only lanthanide that can be affected by redox reactions in low temperature aqueous environments (from more soluble Ce^3+^ to less soluble Ce^4+^)^38^, correlated with Fe (*r*=0.45), Ti (*r*=0.42) and P (*r*=0.38); *p*<0.05 in schist, consistent to their leaching from accessory minerals allanite and xenotime. The results also showed plant-enhanced peaks of La in granite and of Tm in basalt. The effect on La can stem from its electronic structure (no unpaired 4f electrons associated to largest ionic radius) which imprints the lowest ligand complexation potential among REE^24^.

Except for schist, a subtle W-class tetrad effect (concave) overlapped the broader patterns in water (Figure 3B). The most conspicuous are the third and fourth tetrads in basalt, clearly developed under plant and mycorrhiza. W-tetrad depressions, presumably produced due to decreasing strength of REE ion-ligand complexes during mineral-water reactions^39^ have been reported in water-rock interactions and mineralizing hydrothermal fluids^19,40^ and have been described in relationship with a complementary M-class (convex curves) pattern^19,40^ developed during same partitioning event. Our results from erupted igneous rock indicate that ecosystem activity can lead to W-class tetrads during mineral dissolution.

**Figure 4.**
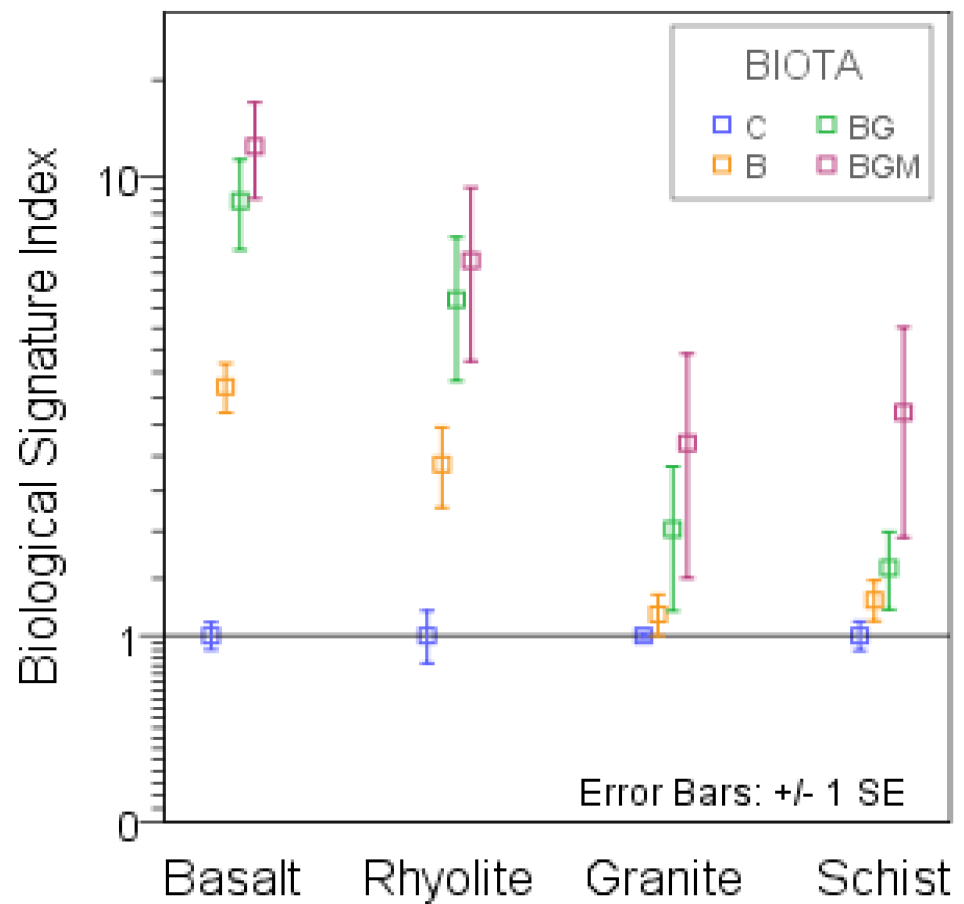
Biotic signature of REE weathering. Lanthanum/phosphate (abiotic-normalized concentrations) in pore water samples collected from the four rock types, averaged across the two-years experiment. The index was calculated by using the Equation s1 and it is unit-less. Treatments: C, control; B, bacteria; BG, grass-bacteria and BGM, grass-bacteria-mycorrhiza. Index values, extending from 1-100 were normalized to the abiotic control of each rock type.

Surprisingly, the factors strongly explaining (*p*<0.05) REE dissolution mechanisms included inorganic carbon forms and host mineral major ions (Table s3). While studies in forested environments have found a strong colloidal carbon control on REE mobilization ^23^, its low influence in this experiment (Table s3) means that during incipient soil genesis, inorganic complexation dominates the mobilization of REE and potentially other metals, this being enhanced by the respiration of organisms. This is also supported by a significant increment (*p*<0.05) in inorganic carbon in water collected from planted treatments. Correlative findings were reported in aquifer studies, where due to less abundant organic ligands in solution, REE complexation seemed dictated by major carbonates and Fe oxy-hydroxides^15,41^. It is, however, expected that organic carbon influence to increase over time, as the ecosystem accumulates more below ground biomass.

**Biological Signature Index (BSI).** In their interaction with nutrient providing substrates organisms modify the mobilized element balances by preferentially dissolution and uptake, and this may be captured as fingerprint in the environment. Results of abiotic-normalized La:phosphate index (Equation s1) showed surprisingly consistent behavior among the studied substrates and biota, with abiotic values in the lowest range followed, in order of ecosystem complexity, by microbial alone, and plant (Figure 4). The strongest signals were in basalt and rhyolite, followed by schist and granite. Phosphates are major REE-bearing minerals, and P is also the ultimate life-limiting element sourced from rock^42^, as it cannot be fixed from the atmosphere. The results are consistent with preferential loss of La-one of the most reactive REE in the cationic series, from its phosphate matrix by complexation with negatively charged biological ligands. This, together with a higher water-normalized La than P found in plants suggests a more soluble La vs P in the presence of biota, hence the recorded signal.

**REE uptake and distribution in plant as affected by rock and mycorrhiza.** REE concentrations in biomass were order of magnitude higher in below-ground as compared to above-ground biomass for most rocks (Figure 5), and they were consistent with values described for natural environment^43^. Plants grown in rhyolite accumulated the most, followed by schist, basalt, and granite. Shoot abundances generally mirrored their root counterparts.

The ratio of shoot to root concentrations followed the order (REE mean±SE): granite (0.95±0.41) ≥ basalt (0.47±0.24) > schist (0.21±0.01) ≥ rhyolite (0.18±0.03). Granite’s high above-ground REE transfer is most likely due to its low uptake (Figure 5). The lower but correlative above-ground abundances are evidence of uptake/transfer controlling mechanisms that work similarly for all rare earth ions. A ring-like cell wall modification in root endodermis (Casparian strip), which protects the xylem from passive (pericellular) diffusion of solutes during water uptake, is thought to restrict the direct transfer of REE to xylem^44,45^ by redirecting the water flux through the selectively permeable plasma membrane of endoderm. This mechanism keeps ions in the more active aerial organs in physiologically-relevant balance regardless of their environmental abundances, thus preventing potentially toxic levels. The cell wall of root cortex (hosting negative charges of carboxyls and hydroxyls) is the most likely structure to retain REE entering the root in our experiment and it has been shown to accumulate REE ions^46^. From our results, the membrane also appears to be more permeable at lower concentrations, e.g. granite’s higher transfer factor, which implies increased restriction at potentially toxic levels, a mechanism that may be common to other heavy elements.

Mycorrhiza increased root uptake in basalt (most REE) and schist (M-and H-REE), and their transfer to shoot (Figure 5). Basalt also had the highest mycorrhiza infection rate, 67±18% (Table s2). In rhyolite, no mycorrhiza effect on REE concentrations in biomass was found despite a 52±30% infection rate and lower water REE abundances (Figure 3), possibly due to larger biomass in this treatment (*p*<0.2; Figure s4)-hence larger total uptake. Plants in granite failed to develop infection (Table s2). The few studies addressing the effect of arbuscular mycorrhiza on plant REE uptake are generally in the context of crop phytotoxicity, and they showed either a decrease^48^ or increase^49^ in L-REE uptake. Our study on natural REE abundances indicates that fungal symbiosis can stimulate phyto-uptake and transfer.

**Figure 5.**
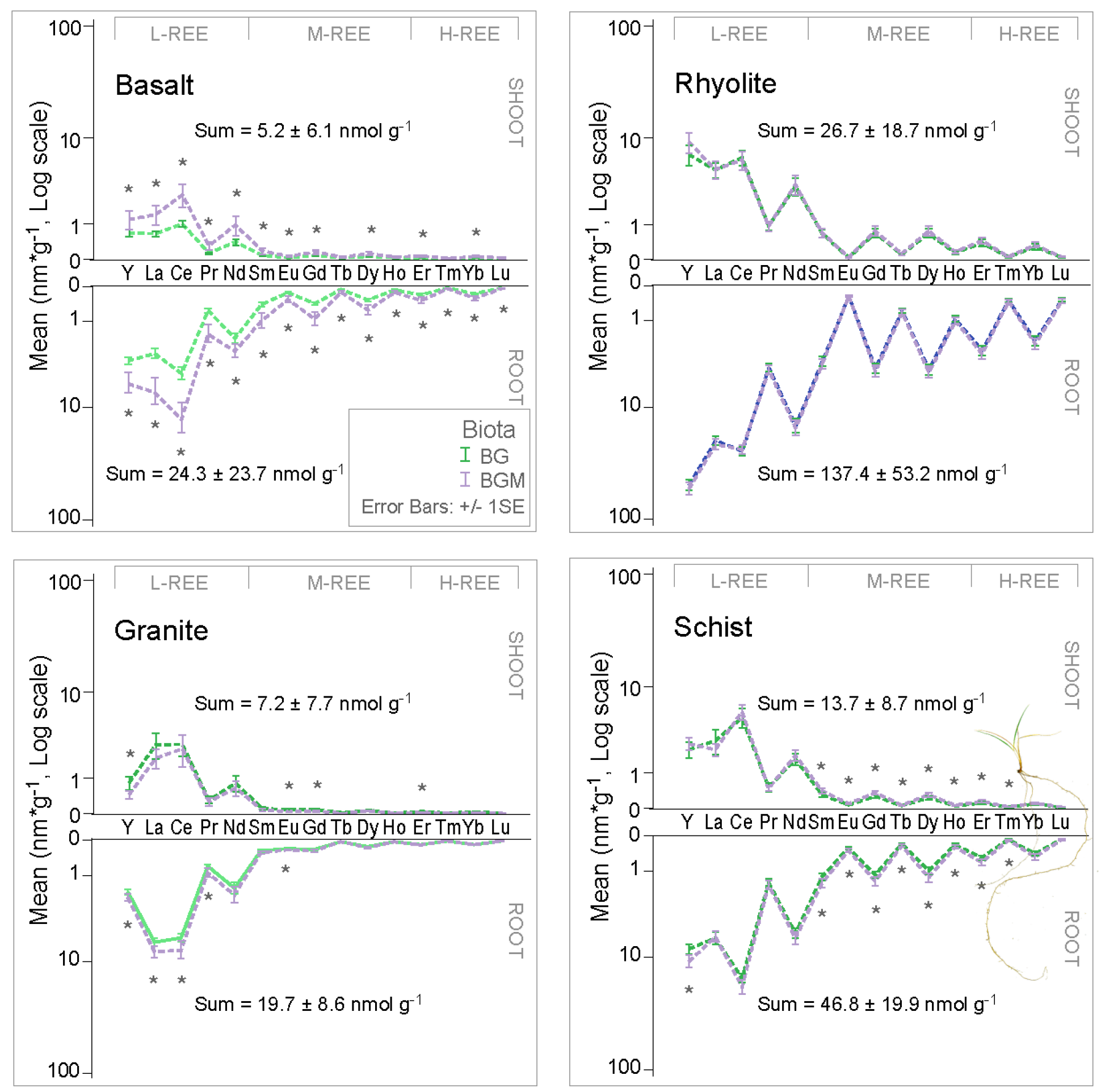
Mycorrhiza effect on REE concentration in plant organs. REE molar concentrations in below (root) and above (shoot) ground biomass of mycorrhiza inoculated and non-inoculated grass growing on the 4 substrates. Significant differences at ±1SE are marked by (*) except for granite, where mycorrhiza infection was unsuccessful.

Water-normalized element abundances in plant (preferential uptake) showed a high uptake efficiency (molar ratio root: water of >1), and supported the concentration findings, lighter elements being favored (Figure 6). This is similar to other vascular plants^43,44^, and can be ascribed to preferential uptake of the more soluble forms, or free ions^50^, owing to lower stability constants of L-REE with inorganic and organic ligands^51^. The shape of the curves generally mirrored their aquatic counterparts (Figure 3), with a subtle M-type (convex) tetrad pattern among H-REE. This indicates that plants were key for the formation of patterns in the water. The tetrads, also found in wheat grown hydroponically with phosphate in the absence of organic ligands^52^ can be produced by the variable adsorption and/or co-precipitation of REE with phosphates and oxyhydroxides on root cell walls^46,53^. This is supported by the REE relationship with P and a limited number of trace (Fe, Sr, Mn, Cu, Ti, Zn, Cr, Al, Ti, and Zn) and major (Na, K, Si and Ca) elements in roots (Table s4). These root processes and patterns also appear to determine REE fractionation in aerial parts (Figure 6). An absence of Eu and Ce anomalies in the plant, means that their dissolution mechanisms have not affected the uptake.

**Figure 6.**
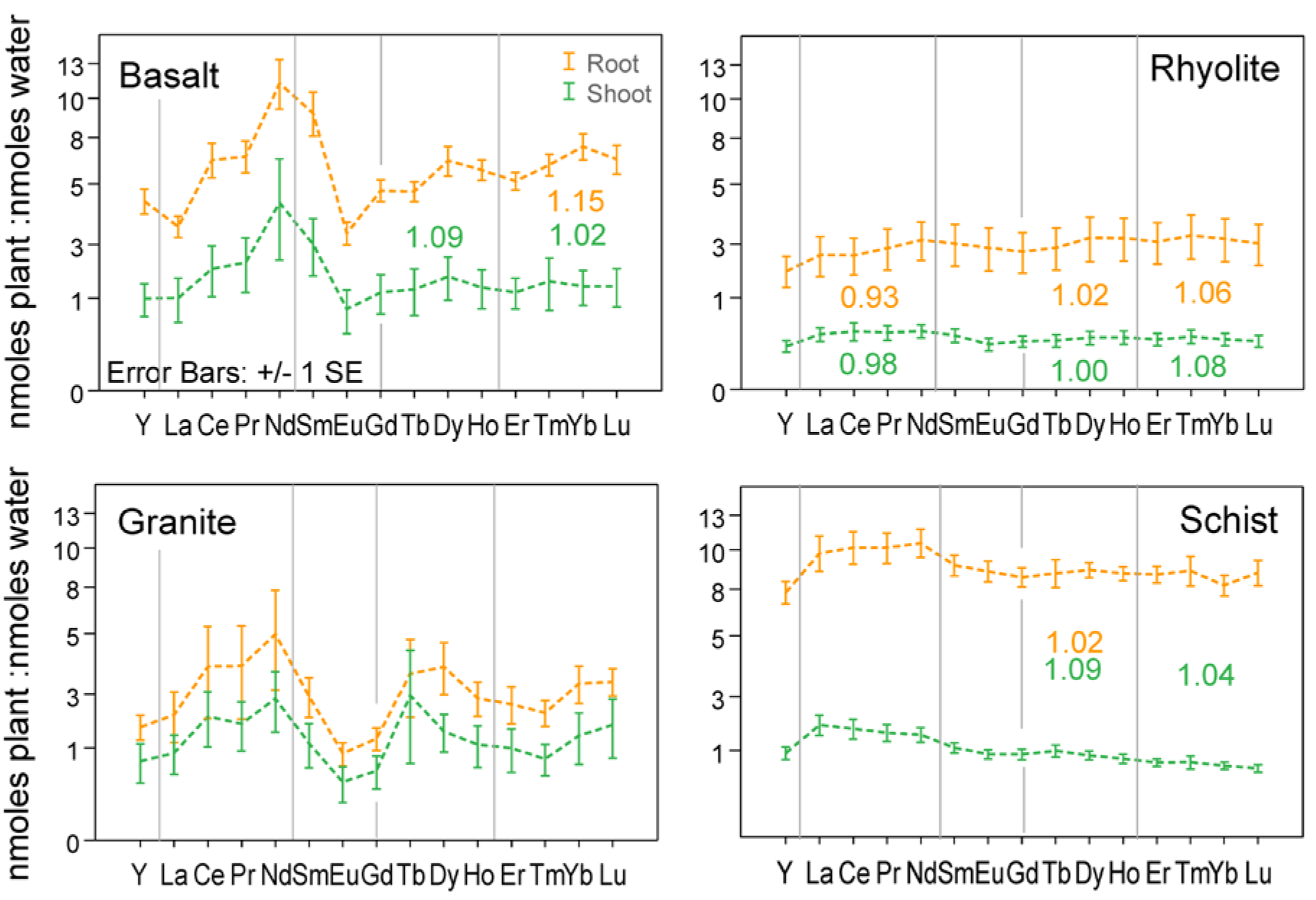
Preferential REE uptake and transfer in plant. Distribution of water normalized REE (total nmoles per column: total nmoles per column, plant:water) in above (shoot) and below (root) ground biomass. Tetrad classes are delimited by vertical lines. Numbers inset represent tetrad magnitude values (numbers range from 1-no tetrad to >1 for M-type and <1 for W-type curves), with color corresponding to treatment color.

**REE in easily available fractions in weathering rock.** Stepwise sequential chemical extraction of water soluble, carbonate and exchangeable (ammonium acetate extractable, AAE) fraction (Figure 7A) showed a biota effect that was opposite to changes in the aquatic phase (Figure 3). Since this fraction can be considered bioavailable, the observed effect can be partially attributed to plant uptake.

**Figure 7.**
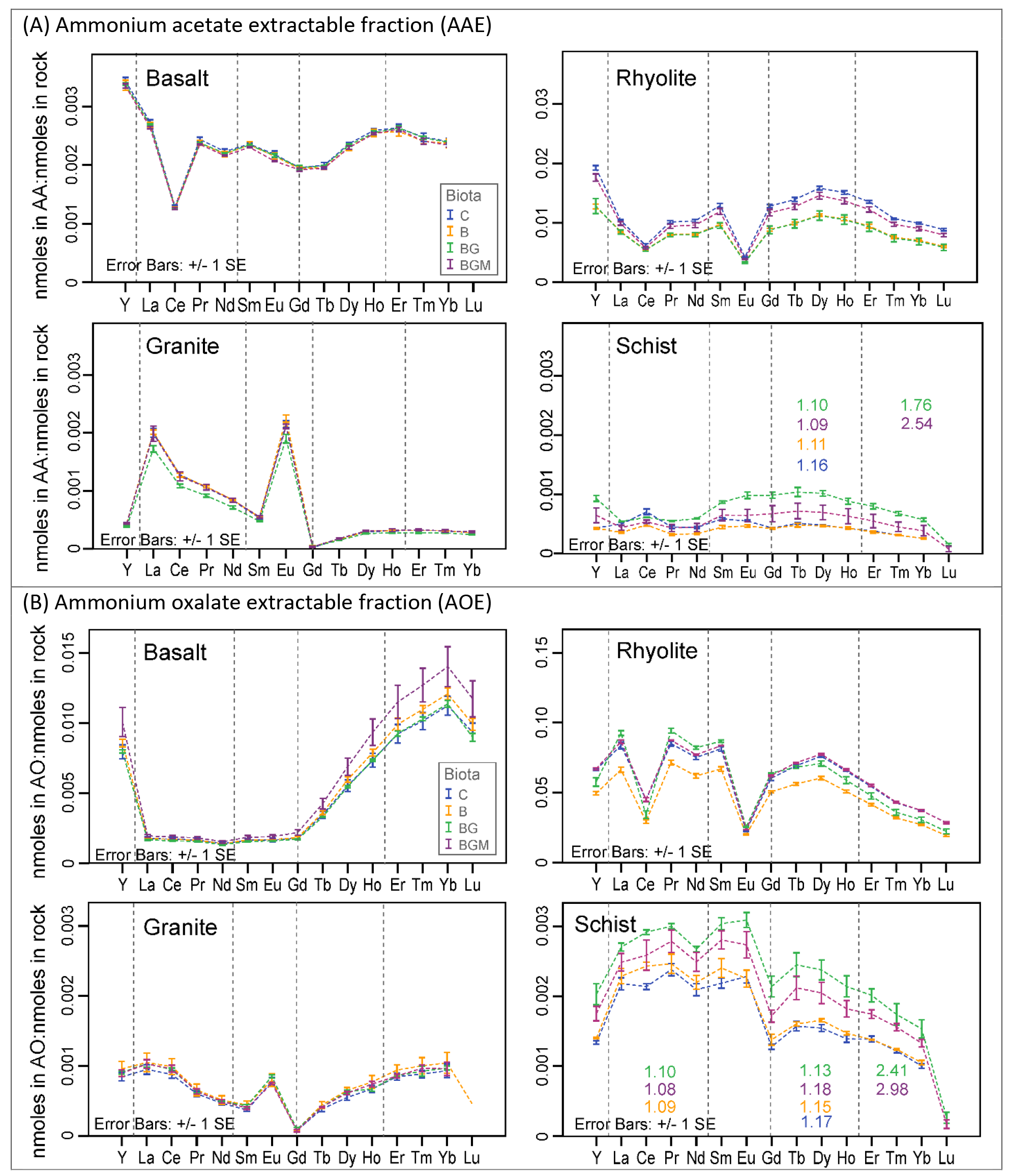
REE variability in easily extractable solid fractions. Rock-normalized REE abundances in (A) ammonium acetate extractable (AAE) and (B) ammonium oxalate extractable fractions of rocks at the end of the 20-months experiment. Values are means of total nmoles/column: total nmoles/column, for triplicate columns. Tetrads are grouped by vertical lines with tetrad value colors representing treatment color.

In rhyolite and granite (both felsic), biota caused a decrease in the AAE pool, particularly in BG treatment, coinciding with a comparatively well-developed plant biomass (Figure s4). The differences were significant at 95% for most elements in rhyolite. In schist, an increase in the exchangeable pool under vascular plant relative to microbial and control (significant at 95% for most REE in BG treatment) coincided with the smallest plant biomass (Figure s4). Mycorrhiza addition increased this fraction as compared to grass-only in rhyolite, and decreased in schist, mirroring the aquatic effect (Figure 3). Results from the first four months of the experiment showed that major nutrient mobilization (denudation and uptake in the vascular plant) is governed by element supply from parent mineral and plant physiological requirements^47^. Therefore, given similar biotic consortia in all rocks, the observed biotic differences in exchangeable fraction among rocks must be directly connected to biological response to major nutrient balances of each rock. Except for basalt and rhyolite, which developed Ce anomaly, probably due to conditions unfavorable for its redox transformation, the general patterns across REE series mimicked their water counterparts (Figure 3).

Analysis of the more stable poorly crystalline (ammonium oxalate extractable, AOE) fraction (including precursors of secondary minerals) revealed that generally, Y and heavier elements preferentially precipitated in basalt, and lighter elements in rhyolite and schist (Figure 7B). A rhyolite Ce anomaly resembling one observed in AAE, as well as Eu anomalies in granite and rhyolite found in water and AAE were also transferred to this fraction. With few exceptions, biotic treatment effects broadly resembled the ones in AAE, and they were largely rock specific. Specifically, microbes significantly (*p*<0.05) decreased REE retention in the poorly crystalline pool (as compared to control) in rhyolite (Figure 7B). The addition of vascular plants increased the retention over microbes in schist and rhyolite. Mycorrhiza increased REE retention into this fraction in basalt (*p*<0.05 for most REE) and it decreased in schist (*p*<0.32). In rhyolite, mycorrhiza generally decreased L-REE and increased H-REE retention (*p*<0.05 for most REE).

An M-type tetrad effect (expected for sediment/soil)^54^ developed in schist in both AAE and AOE fractions (Figure 7). The first tetrad in AOE appeared only in biota systems (Figure 7B). Since our study only covers a short period, it is expected that over longer weathering, or in natural soil genesis setting the amplitude of the effect be greater, and develop on a multitude of substrates. The results of poorly crystalline, exchangeable, and pore water reveal that in newly established ecosystems there is a significant biotic regulation of REE pool as the first step in soil genesis, by promoting REE denudation and stimulating retention in available soils pools, particularly in the vegetated treatments.

**Coupled denudation, uptake, and stabilization in secondary solid phases.** Microbial and plant communities are critical players in elemental balance by mobilizing, accumulating and redistributing elements originating from weathering rocks. Quantitative (mass balance) analysis showed the greatest total REE mobilization (solution and plant) in rhyolite, followed by schist, granite, and basalt (Figure 8). The easily extractable (bioavailable) solid pools ranged from almost 10% of rock total in rhyolite to less than one percent in basalt and tenth of a percent in granite and schist. They were two to three orders of magnitude larger than mobilization and their ratio to mobilization decreased from C to BGM (Table s5). This implies an incremental effect on REE mobilization-sequestration balance with ecosystem complexity in all rocks.

The biological effect on mobilization was significant (*p*<0.05), the difference being caused by both, changes in denudation (dissolved forms), and contribution by uptake. Un-inoculated control, representing baseline abiotic weathering generally experienced the lowest REE denudation, while the microbial community enhanced denudation, with significant effects in rhyolite. This was connected to a decreased exchangeable (AAE) and poorly crystalline (AOE) fractions in this rock, most likely due to the biofilm preventing their formation.

Plant presence significantly increased denudation, from 2.2 times in basalt to 3.0 in granite and 4.6 in rhyolite. As plant biomass presented a greater pool of REE than denudation, the development of vascular plant significantly increased total mobilization in all rocks, which was about an order of magnitude larger compared to inorganic control alone (Figure 8). Most of the plant REE were stored in the root, reflecting a high REE weathering efficiency of root-microbe consortium. In rhyolite and schist, where plant biomass was higher as compared to the other rocks (significant for rhyolite at 67% level), plant induced an increase in poorly crystalline+exchangeable fractions (in both environments). The activity of arbuscular mycorrhiza induced a generalized increased REE mass in plant of 1.2 to 1.6 times (*p*<0.05), but in rhyolite an increased plant uptake was balanced by decreases in dissolved forms (*p*<0.05). As a result, total mobilization was not significantly affected by mycorrhiza in this rock. However, exchangeable and amorphous fractions were incremented under mycorrhiza in rhyolite.

It is also to be noticed that both exchangeable and amorphous pools were relatively large in the initial rock (Figure 8). We assume that this is due to initial high reactivity of fresh mineral surfaces when open crystal lattices resulted from rock grinding easily release elements to chelating agents. However, given that a significant biotic effect was observed in these fraction (Figure 6), it is safe to assume that a portion of it represents novel exchangeable and amorphous phases resulted from weathering. The results suggest that in the transition from simple to complex, e.g. in ecosystem colonization of freshly exposed bedrock including fresh volcanic fields, mountain tops, surfaces exposed by glacier retreat, and soil bedrock horizon, biota has major control of REE cycle initiation. Secondly, since roots of vascular plants have taken up a significant mass of weathered REE, they are good sensors, integrators and regulators of REE cycle in the biosphere.

**Figure 8.**
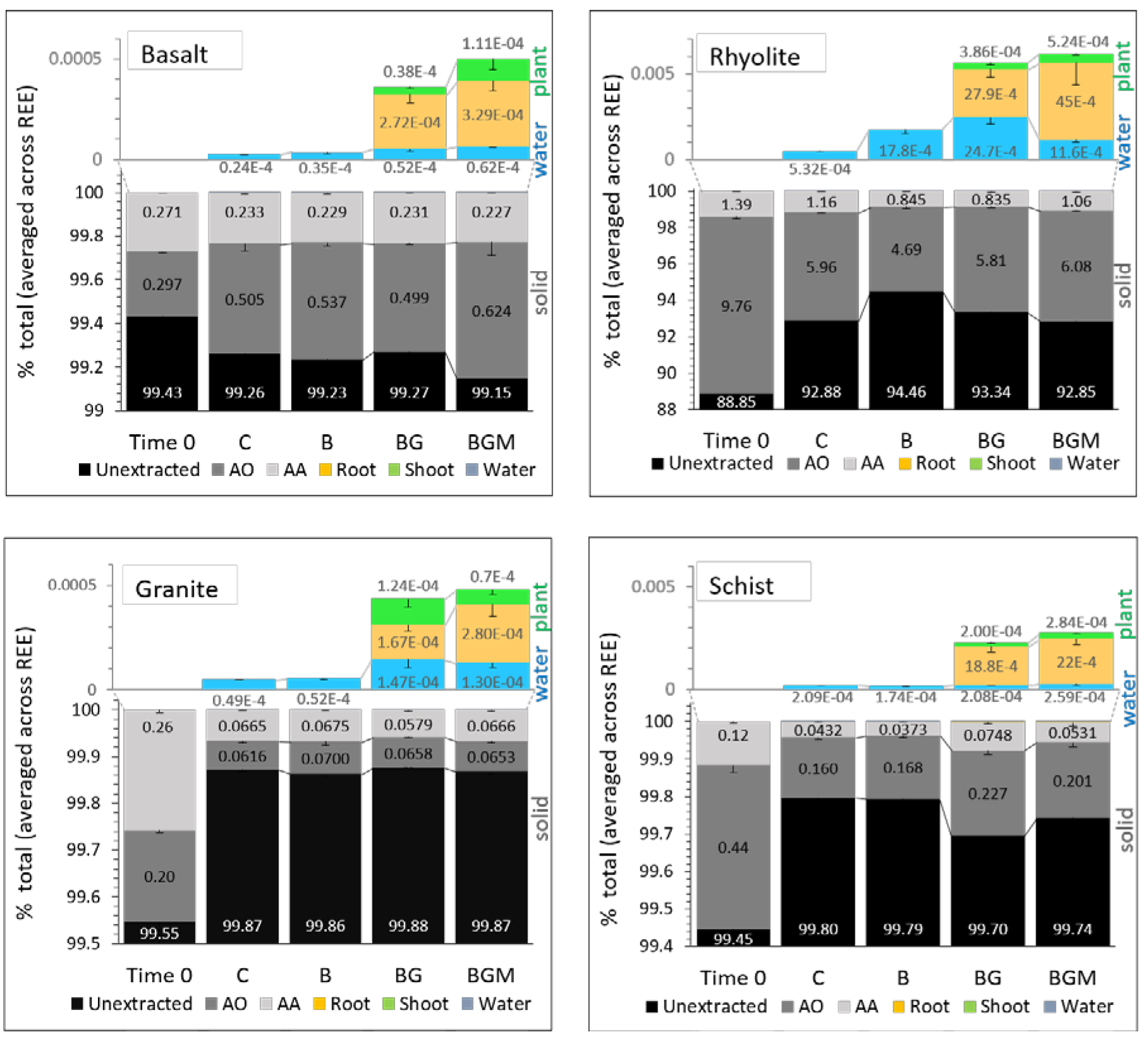
Mass balance of abiotic and biotic REE weathering. Distribution of sum of REE at time=0 and the end of the 20-month experiment as a function of rock type, abiotic and biotic treatments (C, control; B, bacteria; BG, grass-bacteria; BGM, grass-bacteria-mycorrhiza). Shown REE pools include unextracted (black), ammonium oxalate extractable (brown), ammonium acetate extractable (grey), denudation (blue), root biomass (ochre) and shoots biomass (green). Values are means of column triplicates. For each column values were summed across REE series, and for water also across sampling events. Error bars represent +/-1SE.

## Conclusions

The incipient biotic system developed long-term using only nutrients derived from rock. Initial weathering progressed fast, with little difference between biological treatments—mineral dissolution being dominated by abiotic processes. In approximately two months leaching rates declined while microorganisms and plants started to exert greater uptake, dictating REE fate in the ecosystem. Results further indicated that inorganic dissolution enhanced by biotic respiration was predominant throughout the experiment, consistent with low organic matter presence.

The REE mass balance between water, plant and secondary solid fractions (exchangeable and poorly crystalline) varied slightly with rock, but several consistent trends were observed. At the end of the experiment, ten times as much REE amount was mobilized in planted treatments compared to abiotic, with plants hosting a significantly larger REE pool than water alone. Roots stored greater REE mass compared to shoots due to both, greater biomass and greater concentration, meaning that below ground biomass is the main biological store of REE in the ecosystem. Comparatively lower shoot concentrations can be ascribed to REE retention with P and other major ions in root endoderm’s cell walls.

Total REE mobilization (leaching and plant uptake) was generally increased by arbuscular mycorrhiza, but in rhyolite decreased leaching was associated with greater retention in the exchangeable fraction. This type of symbiosis is, therefore, a significant contributor to REE cycle in the ecosystem by controlling REE export and retention balances. Microbes, while having a smaller effect on REE denudation and total mobilization than plants, resulted in significant increases in both parameters in rhyolite in association to lower masses of bioavailable solid fractions, most likely due to the biofilm preventing their formation. The results also imply that more complex ecosystems are generally capable of preferentially mobilizing, accumulating and redistributing higher masses of REE in the environment than simpler ones, even early in the biological colonization. This was also supported by bioavailable solid pools decreasing relative to the dissolved and plant pools as treatment complexity increased.

A synergetic interplay between rock, water, and biota determined variability patterns among REE, with both biotic system and rock mineralogy imprinting differences among these patterns in association with major cations. REE in biota can therefore directly reflect soil genesis processes, including rock weathering/denudation capacity. Moreover, the similarity in REE patterns detected between root and shoot makes above-ground biomass a reliable integrator and sensor of REE signals from belowground ecosystems. There was a strong biotic signature in REE weathering, supported by the weathering magnitude, the general pattern among REE (particularly L-REE), the development of a tetrad effect, a consistently increased REE mobilization-sequestration ratio with ecosystem complexity, and a consistent REE:phosphate BSI index in water in all rocks. This shows that selective fractionation by ecosystem exists and it can drive REE partitioning during bio-weathering, leaving detectable signals of their presence in the environment.

Using current data on global weathering estimates derived from river fluxes and water elemental stoichiometry in the experiment (SI 1.3), we estimated a total annual REE denudation rate of about 3858.88*10^4^ moles from the four exposed lithologies. Of this, 60.45*10^4^ moles are contributed by basalt, 924.13*10^4^ moles by rhyolite, 93 2.66*10^4^ moles by granite and 1941.65*10^4^ moles by schist. Abiotic weathering alone contributed about 38.6±19 % of these values (12.4 basalt, 48.5 rhyolite, 30.4 granite, and 63.1vschist, % of GBM), with the remaining added by the biological component of the ecosystem. Given that solute denudation to oceans is dominated (>50%) by steepest 10% of Earth’s surface, i.e. mountains^55^, it is safe to assume that most REE weathering happens in areas of high weathering but young soil, which is comparable to our experiment. However, these estimates are clearly a first-order approach to understanding global REE fluxes. Our results provide a quantitative understanding of biological contribution to REE weathering in common upper crust rocks, which can have broad implications in modeling their global cycles and budgets in relationship with overlaying ecosystems, as well as for understanding the potential biological signatures accumulated in Earth’s historical record, as well as on other planetary bodies.

## Methodology

**Experimental design and solution analysis.** A model ecosystem experiment was setup in the Desert Biome at University of Arizona’s Biosphere 2 (SI Figure 1). Four granular substrates (basalt, rhyolite, granite and schist) and six ecological treatments were placed in 288 experimental columns (30cm long x 5cm internal diameter) in six controlled-environment chambers receiving filtered and UV light-sterilized air and purified water. For this study, we report the effect of four biological treatments on the weathering of the four bedrock types during the first 20 months of colonization. In increasing order of complexity, treatments were: control (with autoclaved microbial consortium) (labeled as C); microbes (a natural consortium extracted from basalt collected in Merriam crater, Flagstaff, AZ) (labeled as bacteria, B); microbes and grass (Buffalo grass, *Bouteloua dactyloides*) (BG); and microbes, grass and mycorrhiza (grass infecting *Rhizophagus irregularis*) (BGM). Treatments were run in triplicate with water samples collected over 21 time points (December 2012-July 2014). To allow solid and biotic phases analysis over four time points (at 132, 252, 465 and 584 days), treatments were repeated four times.

The substrate materials were collected from Santa Catalina Mountains (medium-grain Oracle granite and micaceous Pinal schist); Meriam crater, Flagstaff, AZ (cinder basalt); and Valles Caldera National Preserve, NM (rhyolite). Studied rocks were part of the Catalina-Jemez Critical Zone Observatory and the Biosphere 2 Landscape Evolution Observatory, two related large-scale experiments that study the coupled chemical, hydrological, and ecological processes that shape the Earth’s surface and support most terrestrial life^2,56^. Except for basalt (fresh tephritic material of very limited weathering, which was ground at the mining site), the rocks were subjected to mechanical removal of all weathered crust, before being crushed and ground. All substrates were then dry sieved and wet sieved to 250-500μm particle size, passed on a Wilfley water table to remove impurities, and rinsed several times with nanopure-grade water. The clean material was then dried at 85°C. Substrate characteristics are in Table s1. The rock was then sterilized by autoclaving for one hour over 3 consecutive days. Sterilization was confirmed by a lack of colony-forming bacterial growth on R2A agar plates.

Each column received 90 mL of inoculum containing 1.43×10^5^ colony forming bacteria per mL under sterile hood, incubated at room temperature for one week in sterilized jars, packed to top of the column in UV-light sterilized acrylic columns (Table s1 for rock weight) and planted with dehusked and pre-sterilized germinated grass seeds (20 per column, 2cm depth; purchased from Western Native Seed, Colorado, USA)^47^. Control samples were inoculated with a sterilized inoculum to retain a consistent chemical elemental composition. A 1-mL water suspension of pure *R. irregularis* spores and mycelium (about 400 spores; Premier Tech Biotechnologies, Canada) were added to each mycorrhizal treatment column next to the seedling.

A sterilized 140mL polypropylene syringe (Nasco, Modesto, CA) was used to add 100-120mL of sterile nano-pure water (18MΩ) per column every two weeks for the first two months and monthly after that, allowing collection of 35-50mL sample. Watering was interpolated to allow plants receive water every two weeks. The irrigation system was designed to prevent preferential water flow in the substrate. Water samples were analyzed for pH, electric conductivity (by electrode), total organic carbon (680°C combustion catalytic oxidation, Shimadzu TOC-L system) and major, trace and REE as rock samples by ICP-MS (Perkin Elmer, Elan DRC-II). Major anions, i.e. fluoride, chloride, nitrite, bromide, nitrate, sulfate and phosphate were analyzed by ion chromatography (Dionex). The water balance was determined by subtracting water output volume from water input at the time of sampling, and it is assumed to be an indication of substrate water retention capacity (retained, transpired and evaporated by biota and column surface).

**Initial rock characterization.** Mineralogical characterization was performed on all rock samples (250-500μm size fraction) by electron microprobe and X-ray diffraction. Microprobe data were collected using CAMECA SX100 Ultra and CAMECA SX50 electron probe microanalyzers (Lunar and Planetary Sciences Laboratory, University of Arizona, USA). A beam size of 5μm (15KeV, 20nA) was chosen to probe chemical composition in feldspars and glass and 1μm for other minerals. For the calculation of chemical formulae, the elemental composition was normalized to oxygen. In addition, 5×5mm elemental maps were collected to characterize element heterogeneity of the sample, rock microstructure, and to provide additional means of determining mineral composition. High current mode (25KeV and 20nA) was used for identification and mapping of selected REE and their mineral hosts.

To quantify mineral abundance, samples were micronized to <2μm and subjected to high energy X-ray diffraction (XRD) on beamline 11.3 at the Stanford Synchrotron Radiation Lightsource (SSRL), USA. The line was operated in transmission mode at ca. 12735eV, using a 34.5cm radius Mar detector with 100μm pixels. Three scans were collected for each 0.05g sample and combined. Quantitative analysis of minerals was performed using the Rietveld module included in the X’Pert HighScore Plus software. Total elemental composition of rock samples was measured following lithium/tetraborate fusion followed by inductively coupled plasma-mass spectrometry (ICP-MS, Activation Laboratories Inc., Ancaster, Ontario, Canada) analysis.

**Plant material analysis.** After 20 months, columns were sealed under sterile conditions, extracted from growth chambers, and stored at 4C° until lab processing. Under a laminar-flow hood, plants were separated from loosely attached rock, washed with nanopure-grade water to remove remaining particles, dried at 70°C for 3 days, separated into above and below ground biomass, and weighed. Subsamples of above and below ground biomass were microwave digested in 1:1 70% HNO_3_-30% H_2_O_2_ mixture, at 200°C. REE concentrations were measured by ICP-MS together with the same suite of major and trace elements. Standard quality control measures including sample blanks, certified reference material (apple leaves CRM 1515) and triplicate sample analysis were implemented for each digestion batch.

**Sequential extraction.** Following removal from the column, bulk sand was air dried, homogenized, subsampled, and subjected to a two-step sequential chemical extraction following a modified protocol from (^57^). The procedure extracted (a) soluble, exchangeable ions and carbonates (using 0.2M ammonium acetate adjusted to pH 4.5) and (b) amorphous-to-poorly crystalline fraction (using 0.2M ammonium oxalate adjusted to pH 3.0 with 0.2M oxalic acid).

**Data analysis.** To understand the broader context of substrates used, REE abundances in the 4 rocks were analyzed together with their terrestrial upper continental crust values^32^, mantle and carbonaceous chondrite (protoplanetary material)^58^ by Principal Component Analysis (PCA). PCA is a statistical method that can reduce a great number of variables to few composite variables (principal components) that can represent major trends associated with the dataset. Rock concentrations were then normalized by upper continental crust average values to identify potential REE enrichment/depletion. PCA was further used to identify associations between rock-normalized REE, major and trace element abundances in plants, and between REE and major elements in sequentially extracted phases.

ANOVA with Fisher’s least significant difference (LSD) posthoc test of inter-treatment comparisons was used to test the effect of mycorrhiza on plant REE uptake, and the effect of various ecosystem components on total REE mass distribution among weathering pools in a mass balance analysis. Preferential uptake was assessed by dividing total element in the plant (mass) to total removal/ leaching in water (mass).

Tetrad effect was identified by visual inspection of tetrad shapes in bulk rock (water)-normalized concentrations in water, plant and sequential extraction materials. The effect size (t) was determined for each of the four tetrads (i) using the Eq. (1)^40^, which is the equivalent of the method proposed by (^59^).

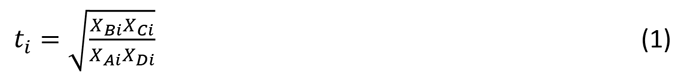

Where *X_Ai_* and *X_Di_* are the normalized concentrations of first and last tetrad members, and *X_Bi_*-*X_Ci_* is the normalized concentrations of the central elements. The formula compares the center members with an imaginary line that connects the first and last members in a logarithmic plot. The values of ti are larger than one for convex (M-class) curves, equal to one for a straight line, and <1 for concave (W-class) curves. Tetrads displaying Ce and Eu anomalies were not quantified. Due to a typically high intrinsic (measurement) error in REE quantification, inter-treatment comparisons of tetrad effect were on ±1 standard error (SE). Statistical analyses were computed in IBM-SPSS statistics v.21.

## Acknowledgements

This research was funded by National Science Foundation (NSF) grant EAR-1023215 “ETBC: Plant-microbe-mineral interaction as a driver for rock weathering and chemical denudation”. We also acknowledge additional support from NSF EAR-0724958 and EAR-1331408 grants that support the Catalina – Jemez Critical Zone Observatory (CZO), the Biosphere 2 REU program, NSF EAR-1263251 and NSF EAR-1004353 (http://www.b2science.org/outreach/reu), United States-Mexico Commission for Educational and Cultural Exchange (COMEXUS): the Fulbright-Garcia Robles Scholarship program; and Thomas R. Brown Foundation endowment to University of Arizona.

We are deeply thankful to Nicolas Perdrial, Julia Perdrial, Nate Abramson, Juliana Gil Loiaza, Joost Van Haren, Peter Troch, Miranda Galey, Elise Munoz, Jake Kelly, Vanessa Yubeta, Lauren Guthridge, Mathew Clark, James Olmid, Guillermo Molano, Andrew Toriello, Nicolas Sertillanges, Arturo Jacobo, Yadi Wang, Julie Neilson and the multiple other collaborators for their field, lab and theoretical contribution.

## Authors Contribution

The corresponding author DGZ prepared the manuscript and supplementary information, all figures and tables, and was the leading researcher of mesocosm design and setup, rock collection and preparation, sample gathering and data analysis. Coauthors KD, JC, RMM, and TH led the grant design and steered its execution. Coauthors CIB and JKP were deeply involved in mesocosm and experimental setup. Coauthors CIB, JKP, EEG, MAPM and MOVI contributed to water collection and plant harvest and digestions, dilutions, as well as data management. Coauthors RMM, EAH, and MKA contributed lab support. Coauthors EAH and MKA analyzed water, plant and sequentially extracted fractions on the instruments. Coauthor KJD contributed electron microprobe support and analysis. DGZ integrated XRD and microprobe data. Coauthors JC, KD, RMM, EEG, JKP, and MOVI were deeply involved in manuscript review.

## Additional Information

**Competing financial interests**: The authors declare no competing financial interests.

## Supplemental Information

### SI 1 Supplementary Methodology

#### SI 1.1 The experiment

**Figure s1.**
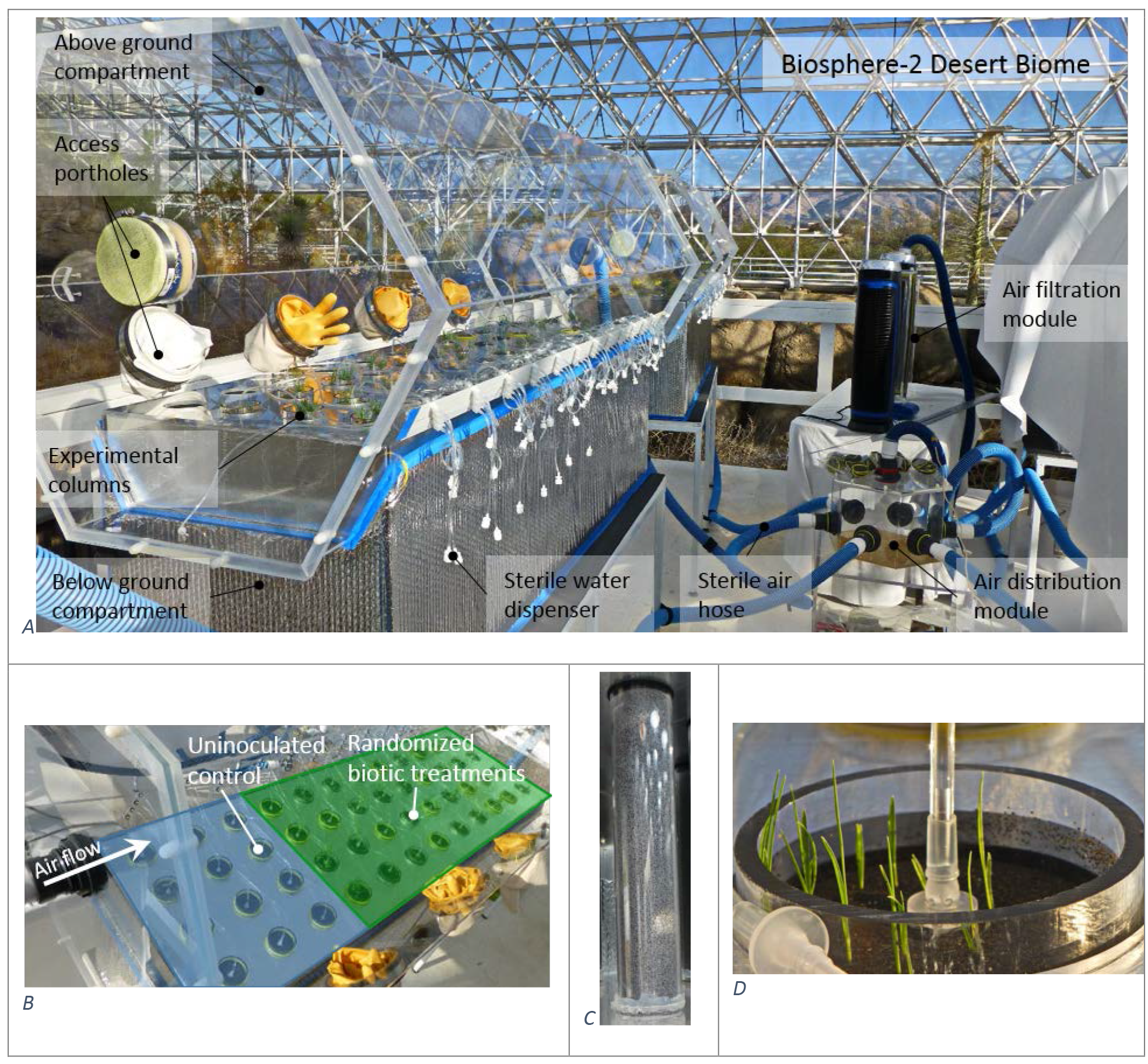
Experimental mesocosms in the Desert Biome of Biosphere-2. (A) containing granular rock-filled columns and purified air and water delivery systems. (B) Treatment distribution in the modules with uninoculated control first in the direction of air flow. (C) Example of experimental column filled with granular granite. (D) Two week old Buffalo grass growing in granular basalt, with water dispenser designed to avoid preferential water percolation in the substrate.

A model ecosystem experiment was setup in the Desert Biome at University of Arizona’s Biosphere-2 based on a design detailed in (^1,2^). Briefly, 6 enclosed chambers connected in parallel to a double air purification system (using 2 high-efficiency particulate absorption HEPA filters and 2 UV-B air sterilization light sources, capable of delivering about 1L air sec^-1^ per module; Germguardian, AC4850CAPT Digital 3-in-1 Hepa Air Purifier System) hold 288 experimental columns (30 × 5 cm internal diameter; Figure s1). Except for control columns, which were placed first in the direction of air flow, the columns were grouped by rock type and randomly distributed in each module.

#### SI 1.2 Biological signature index

To infer a biological signature index, the following lines of evidence were considered: (a) REE sources in phosphate minerals, REE-oxides, and minor minerals (ilmenite, titanite, zircon, allanite) in the used rocks; (b) REE mobilization under biotic treatment was radius dependent, L-REE exhibiting increased mobilization under biota; and, (c) P is the mineral constituent of principal biotic relevance. Based on these premises we propose using an abiotic control-normalized ratio of La to phosphate water concentrations as biotic signature index, following the equation *s1*.

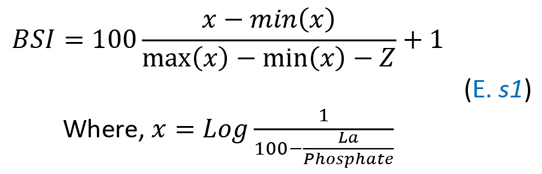

Where, the term 100(*x*-min(*x*))/(max(*x*)-min(*x*) represents a 0-100 scale normalization. *X* is a Log-normalization of La:Phosphate, and Z, a rock-specific constant, representing the abiotic fraction of x. Based on water data of our control treatment Z takes four values (mean±SE): Z_Basalt_=0.543±0.124; Z_Rhyolite_ =1.53±0.339; Z_Granite_=0.626±0.384; and, Z_Schist_=1.23±0.315.

#### SI 1.3 Global denudation estimates

Estimated values for global (G) REE denudation rates were inferred by stoichiometric adjustment of Na-normalized total REE in our experiment (i) to global Na values from river data^3^, according to equation *s2*.

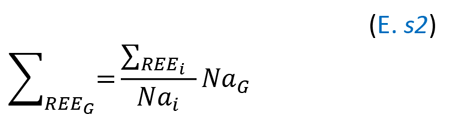

For each rock type, its global Na values were adjusted to the relative contribution of the rock to the global exposed lithology described in (^4^).

### SI 2 Supplementary Results

#### SI 2.1 Substrate characterization

##### SI 2.1.1 *Rocks chemistry in the global context*

Multivariate analysis of rock REE abundances showed greater similarity of the used substrates with the upper (basalt, rhyolite and granite) and lower (schist) terrestrial crust (> 80%variability in dataset), than with the upper mantle and protoplanetary material (carbonaceous chondrite) (Figure s2). Therefore, REE abundances in the substrates were not exceptions to average values of terrestrial crust.

**Figure s2.**
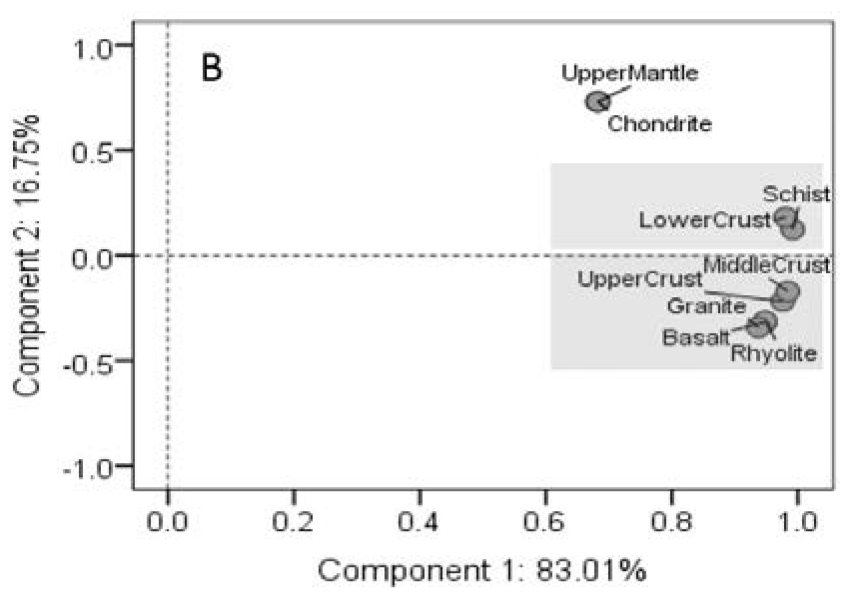
Study rocks in the context of terrestrial and protoplanetary material. Samples were associated based on their similarity in REE content by Principal Component Analysis. Shaded in grey are data strongly associated to component 1 that explains most of the variability in the dataset.

##### SI 2.1.2 *REE mineral source*

**Figure s3.**
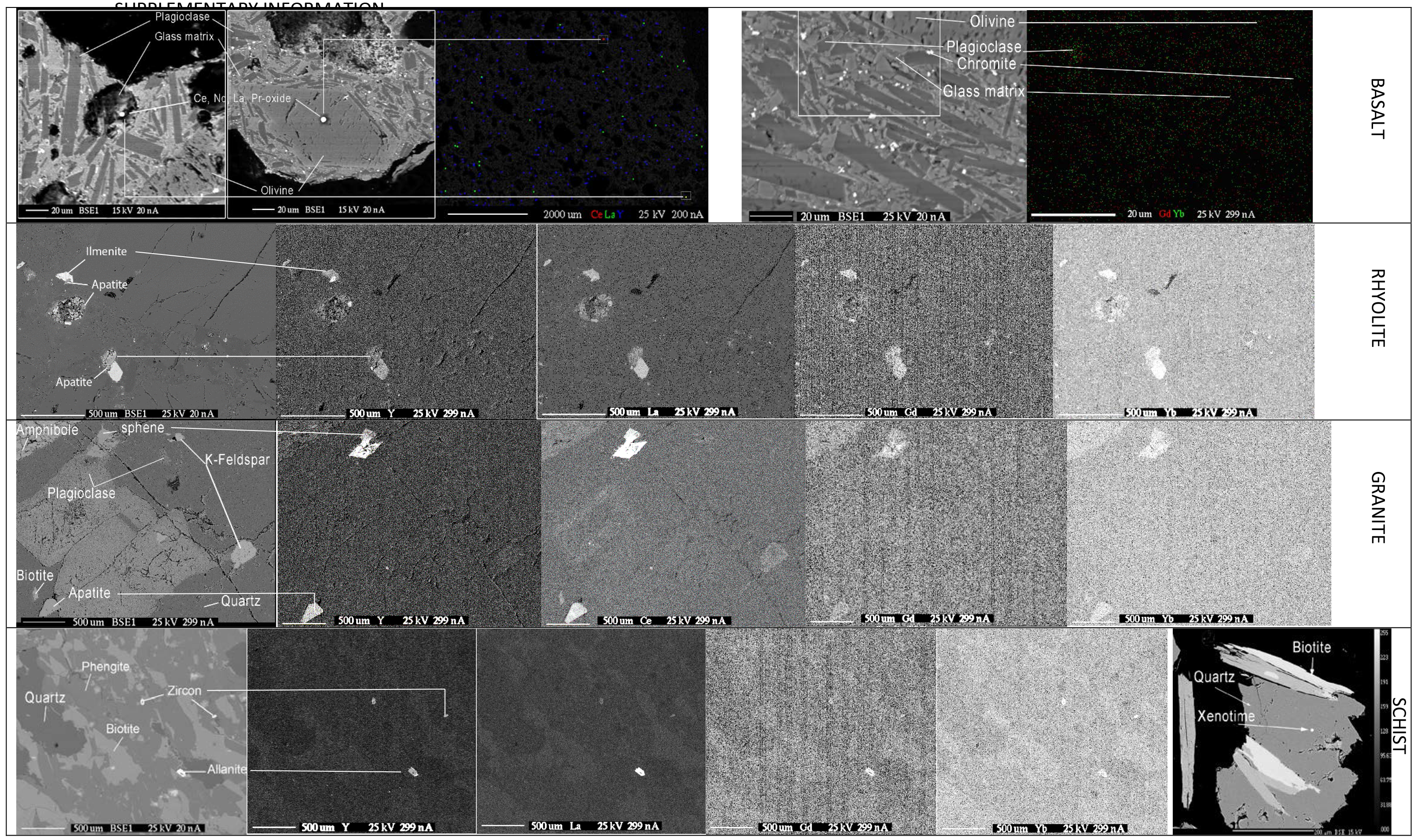
REE mineral source. Back scatter images of studied rocks and high current electron microprobe maps of representative L-, M-and H-REE in the studied materials, and the limited number of minerals that host them. Scattered X-rays appear brighter with increasing REE density. Ce-Nd-La-Pr oxide in basalt, and allanite and xenotime in schist have been identified as main REE mineral hosts. Pixel size in multielement maps has been exaggerated 4X to improve perception of otherwise very low levels.

##### SI 2.1.3 *Substrate physical and mineral characteristics*

**Table s1.** Substrate characteristics (expanded from [^2^]). Mineral formulas were calculated from electron microprobe elemental analyses (a mean of several point analyses), and mineral abundances were estimated quantitatively by Rietveld analysis of X-ray diffraction data. Table headers contain additional information on physical properties of substrates used in the experiment.

| **Basalt** | 690 ± 6 g column<sup>-1</sup> | Density: 1.32 ± 0.03 g cm<sup>-3</sup> | Pore volume: 33 ± 0.7% | **Average %** |
| --- | --- | --- | --- | --- |
| Volcanic glass | Ca<sub>0.36</sub>Mg<sub>0.22</sub>Na<sub>0.16</sub>K<sub>0.09</sub>Mn<sub>0.01</sub>Fe<sub>0.49</sub>Al<sub>0.58</sub>Ti<sub>0.11</sub>(HPO<sub>4</sub>)<sub>0.05</sub>Si<sub>1.87</sub>O<sub>5.89</sub> |  |  | NA |
| Andesine |  |  |  | 50.5 |
| Augite | Ca<sub>0.86</sub>Na<sub>0.03</sub>Mg<sub>0.78</sub>Fe<sup>2+</sup><sub>0.27</sub>Al<sub>0.22</sub>Ti<sub>0.05</sub>Si<sub>1.81</sub>O<sub>6</sub> |  |  | 21.8 |
| Forsterite | Mg<sub>1.55</sub>Fe<sub>0.35</sub>(Si<sub>1.04</sub>O<sub>4</sub>) |  |  | 21.2 |
| Labradorite | Na<sub>0.32</sub>Ca<sub>0.68</sub>Al<sub>1.56</sub>Si<sub>2.35</sub>O<sub>8</sub> |  |  | 3 |
| Titanomagnetite | Fe<sub>2.19</sub>Ti<sub>0.55</sub>Mg<sub>0.21</sub>Al<sub>0.15</sub>Si<sub>0.11</sub>O<sub>4</sub> |  |  | 2 |
| Minor minerals: | Chromite (* (Fe<sup>2+</sup><sub>0.66</sub>Al<sub>1.24</sub>Mg<sub>0.64</sub>)Cr<sub>0.51</sub>O<sub>4</sub>), Apatite |  |  | < 1 each |
| **Rhyolite** | 690 ± 10 g column<sup>-1</sup> | Density: 1.24 ± 0.02 g cm<sup>-3</sup> | Pore volume: 35 ± 1% |  |
| Oligoclase | Na<sub>0.69</sub>Ca<sub>0.24</sub>Al<sub>1.23</sub>Si<sub>2.76</sub>O<sub>8</sub> |  |  | 36.7 |
| Sanidine | K<sub>0.36</sub>Na<sub>0.54</sub>Al<sub>1.02</sub>Si<sub>2.99</sub>O<sub>8</sub> |  |  | 22.8 |
| Quartz | SiO<sub>2</sub> |  |  | 17.2 |
| Anorthoclase | Na<sub>0.70</sub>K<sub>0.23</sub>Al<sub>1.05</sub>Si<sub>2.94</sub>O<sub>8</sub> |  |  | 16.7 |
| Ilmenite | Fe<sup>2+</sup><sub>0.82</sub>Ti<sub>1.02</sub>O<sub>3</sub> |  |  | 5.0 |
| Titanite |  |  |  | 1.5 |
| Minor minerals: | Titanohematite (Fe<sup>3+</sup><sub>2.49-2.52</sub>Ti<sub>0.1-0.15</sub>O<sub>3</sub>), Faujasite, Phengite, Apatite, Zircon |  |  | < 1 each |
| **Granite** | 640 ± 8 g column<sup>-1</sup> | Density: 1.34 ± 0.003 g cm<sup>-3</sup> | Pore volume: 32 ± 0.8% |  |
| Albite | Na<sub>0.95</sub>Ca<sub>0.003</sub>Al<sub>1.02</sub>Si<sub>3</sub>O<sub>8</sub> |  |  | 32.3 |
| Sanidine | K<sub>0.86</sub>Na<sub>0.15</sub>Al<sub>0.98</sub>Si<sub>3</sub>O<sub>8</sub> |  |  | 18.5 |
| Oligoclase | Na<sub>0.73</sub>Ca<sub>0.24</sub>Al<sub>1.18</sub>Si<sub>2.78</sub>O<sub>8</sub> |  |  | 18.4 |
| Quartz | SiO<sub>2</sub> |  |  | 18.0 |
| Biotite | K<sub>0.91</sub>Mg<sub>1.56</sub>Fe<sup>2+</sup><sub>1.06</sub>Al<sub>1.15</sub>Si<sub>2.83</sub>O<sub>10</sub>(OH)<sub>1.75</sub>F<sub>0.25</sub> |  |  | 6.6 |
| Titanite | Ca<sub>1.04</sub>Ti<sub>0.77</sub>Al<sub>0.25</sub>Fe<sup>3+</sup><sub>0.04</sub>SiO<sub>5</sub> |  |  | 3.7 |
| Apatite | Ca<sub>5.22</sub>(PO<sub>4</sub>)<sub>3</sub>F<sub>0.23</sub> |  |  | 2.3 |
| Minor minerals: | Muscovite (K<sub>1.24</sub>Al<sub>1.34</sub>Si<sub>4.14</sub>O<sub>10</sub>(OH)<sub>2</sub>), Chlorite (Mg<sub>2.73</sub>Fe<sup>2+</sup><sub>1.84</sub>Si<sub>2.96</sub>Al<sub>2.25</sub>Mn<sub>0.12</sub>O<sub>10</sub>(OH)<sub>8</sub>), Ilmenite (Fe<sup>2+</sup><sub>0.74</sub>Ti<sub>0.96</sub>Mn<sub>0.32</sub>O<sub>3</sub>), Garnet (Ca<sub>2.38</sub>Fe<sup>3+</sup><sub>0.83</sub>Ti<sub>1.59</sub>Al<sub>0.63</sub>Si<sub>2.33</sub>O<sub>12</sub>), Magnetite |  |  | < 1 each |
| **Schist** | 480 ± 5 g column<sup>-1</sup> | Density: 1.04 ± 0.02 g cm<sup>-3</sup> | Pore volume: 43 ± 1% |  |
| Quartz | SiO<sub>2</sub> |  |  | 47.9 |
| Muscovite |  |  |  | 21.8 |
| Phengite | K<sub>0.87</sub>Al<sub>2.47</sub>Fe<sub>0.21</sub>Mg<sub>0.18</sub>(OH)<sub>2</sub>Si<sub>3.20</sub>O<sub>10</sub> |  |  | 19.7 |
| Biotite | K<sub>0.94</sub>Mg<sub>1.66</sub>Fe<sup>2+</sup><sub>0.88</sub>Al<sub>1.40</sub>Si<sub>2.87</sub>O<sub>10</sub>(OH)<sub>1.57</sub>F<sub>0.43</sub> |  |  | 10.1 |
| Minor minerals: | Xenotime, Zircon, Allanite |  |  | < 1 each |

Basalt consisted of amorphous glass matrix incorporating andesine, olivine and pyroxene, as dominant phases, with traces of other minerals (Table s1). It is higher in Ca, Mg and Fe, and lower in Si than other studied rocks. The vesicular structure of basalt suggested comparatively faster weathering potential. Rhyolite was rich in feldspars and quartz (Table s1). The geochemistry of rhyolite was similar to granite, but it was richer in Si and Na and had less Ca (^2^). Schist contained a high proportion of Mg-rich phengite, a transitional phase between muscovite and caledonite.

##### SI 2.2 *Major pore-water descriptors*

**Table s2.**
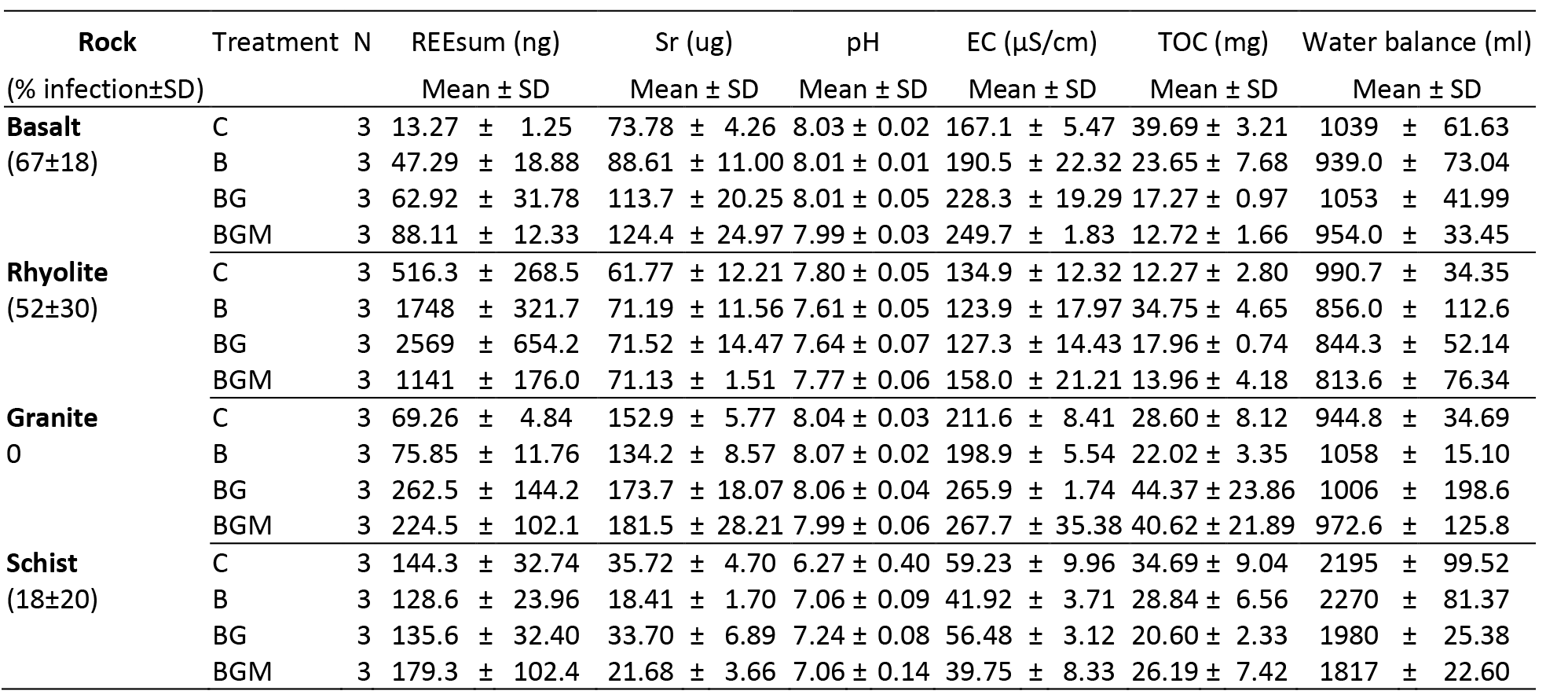
Total REE, Sr (major weathering indicator, for comparison), total organic carbon (TOC), water balance (expressed as difference between input and output volumes) and mean measured pH and electrical conductivity (EC) over the 20-months experiment. Mycorrhiza infection rates are also presented for each substrate.

### SI 2.3 Co-dissolution and uptake mechanisms

#### SI 2.3.1 *Water*

There was a strong correlation among concentrations of different REE in pore water (Table s3) consistent to their group behavior.

**Table s3.**
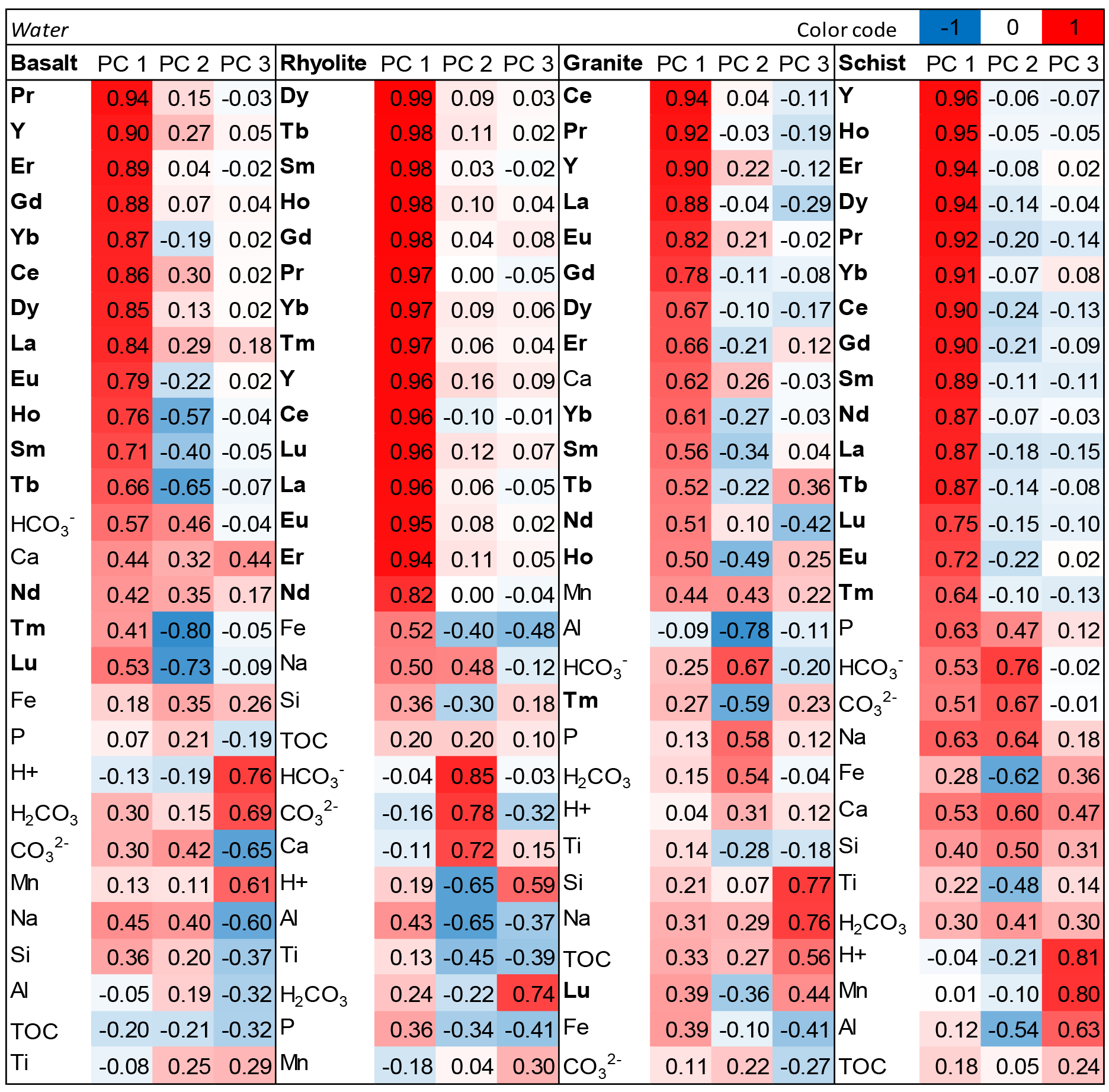
Principal Component Analysis heat map showing correlations between water concentrations of REE, major elements of their mineral sources, total organic carbon (TOC) and other major constituents, that can explain their dissolution mechanisms. Values are ordered in decreasing order of correlation with each component.For easy identification REE are in bold.

#### SI 2.3.2 *Plant*

**Table s4.**
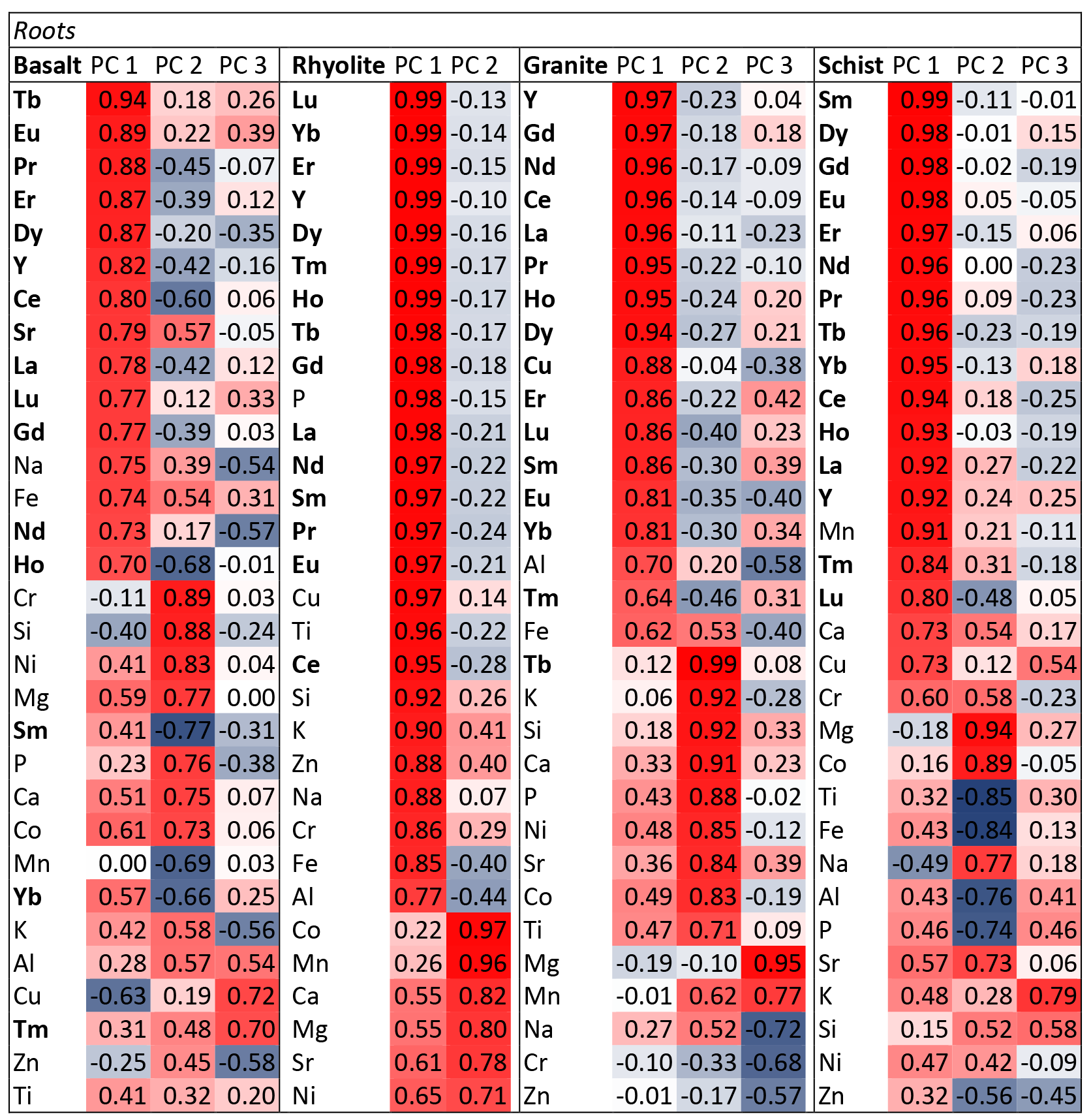
Principal Component Analysis (unrotated) showing relationships between water-normalized REE, trace and major elements in *Buchloe dactyloides* grass roots in different substrates. Correlation values between element and each PC are on a color scale from 1 (red) to 0 (white) and −1 (blue). REE are in bold for easy identification. Elements are in order of their correlation with each component.
SI 2.4 Plant biomass.

### SI 2.4 Plant biomass

**Figure s4.**
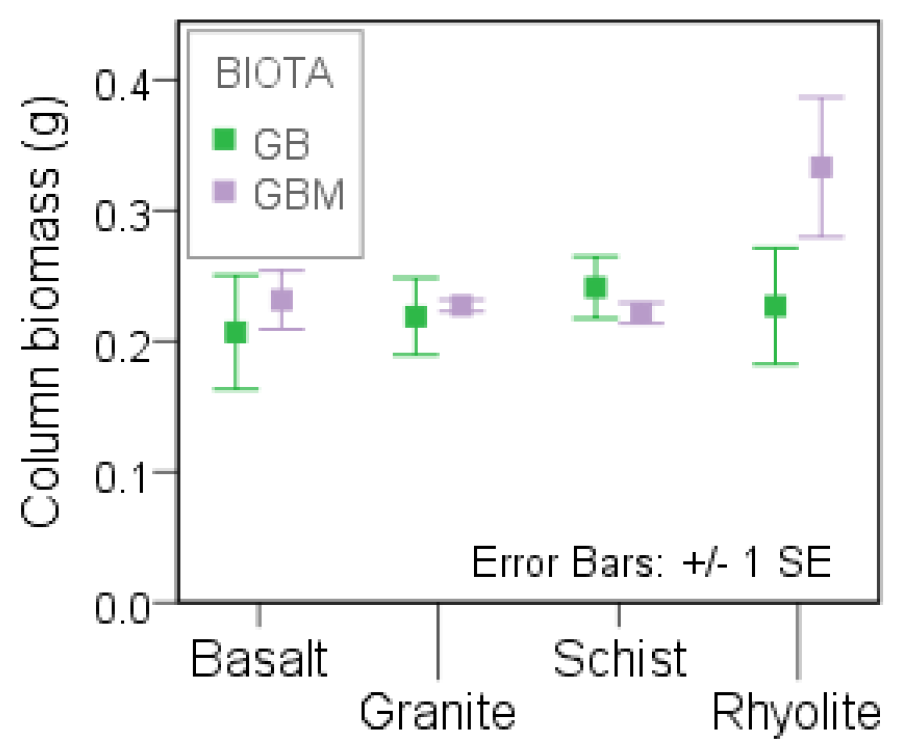
Average column biomass (*Bouteloua dactyloides*) developed on the four rock substrates after 20 months experiment, as influenced by arbuscular mycorrhiza.

### SI 2.5 Coupled denudation, uptake, and stabilization in secondary solid phases

**Table s5.**
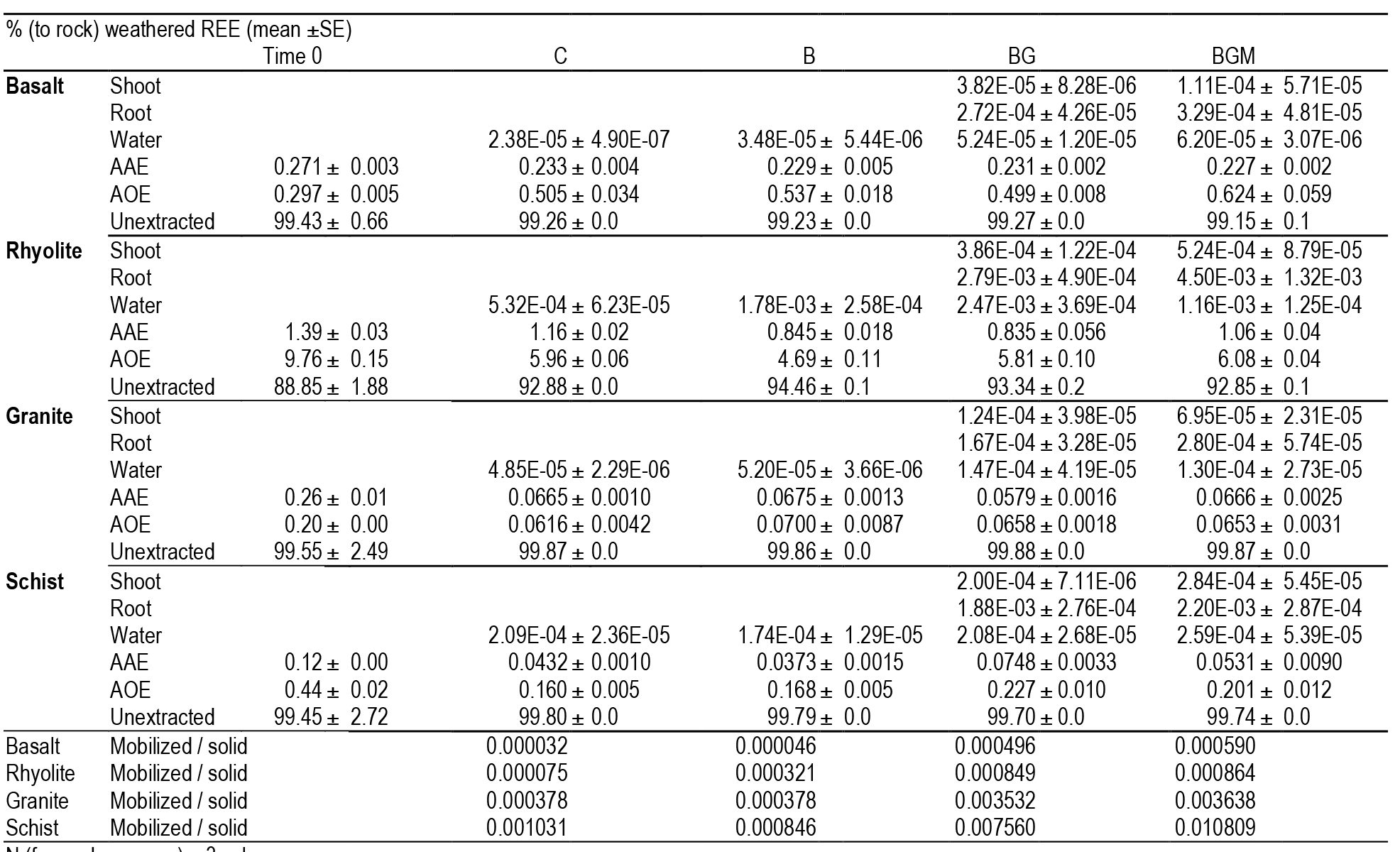
Distribution of REE (sum) in unreacted rock (time=0) and the end of the 20-month experiment for each rock and biotic treatments, together with their ratios mobilized (water + plant) *to* solid phases (AAE + AOE). For each column values were summed across REE series, and for water also across sampling events. Error bars represent +/-1SE. C, abiotic control; B, bacteria; BG, grass-bacteria; BGM, grass-bacteria-mycorrhiza.

